# The first comprehensive case study of early-emerging prosopometamorphopsia

**DOI:** 10.64898/2025.12.15.690312

**Authors:** Sydney Fortner, Antônio Mello, Daniel Stehr, Chandana Kodiweera, Heejung Jung, Krzysztof Bujarski, Tor Wager, Brad Duchaine

## Abstract

Prosopometamorphopsia (PMO) is a rare perceptual disorder characterized by facial distortions. This report presents the first thorough study of a case of PMO that emerged early in life. Zed has experienced lifelong, dynamic facial distortions in which features move, droop, disappear, expand, shrink, and rotate. He also experiences identity misrecognitions, in which he perceives familiar faces as strangers and vice versa (e.g., seeing his grandfather when viewing a young woman). Behavioral assessments indicate that Zed’s distortions occur across viewpoints, visual field positions, picture-plane orientations, and visual angles. Neuropsychological testing revealed face identity recognition deficits, yet, despite his facial distortions, Zed can make accurate fine-grained perceptual judgments about facial age and sex. Structural MRI showed no lesions, but face-selectivity in two right posterior face-selective areas was reduced, and diffusion tensor imaging revealed lower white matter integrity in the left inferior fronto-occipital fasciculus. Zed’s case offers new insights into early-emerging PMO, its manifestations, co-occurrence with other face-processing disorders, and underlying neural correlates.

## 1 Introduction

Prosopometamorphopsia (PMO) is a rare visual disorder characterized by distortions in facial appearance, including alterations in shape, texture, feature positioning, and color. These distortions can affect both sides of the face (bilateral PMO) or just one side (hemi-PMO). In some cases, facial distortions are dynamic, while in others, they remain stable over time ^[1,2]^.

The first documented case of PMO was published in 1904 when Lachmund described a 37-year-old woman, W., who, following three consecutive epileptic seizures, perceived her eyes enlarged and “contorted”, with her head expanding when looking in the mirror ^[3]^. The term “prosopometamorphopsia” was introduced decades later by MacDonald Critchley ^[4]^, who coined the term by combining the Greek prosopon (“face”) with metamorphopsia (the medical term for visual distortion). Since Lachmund’s report, fewer than 100 PMO cases have been documented ^[1,2]^. Nearly all reports involve individuals who acquired PMO later in life, after years of typical face perception. Common causes of acquired PMO include severe head trauma, epilepsy, brain tumors, stroke, and multiple sclerosis ^[1]^. Lesions within the face-processing network are commonly observed in acquired PMO ^[1,2]^, and intracranial stimulation of face-selective brain regions can induce distortions resembling PMO symptoms ^[5–8]^.

In many neuropsychological conditions, however, a substantial proportion of affected individuals exhibit no clear history of brain lesions and are instead classified as having a developmental disorder ^[9–11]^. To date, only one reported PMO case appears developmental in origin. This fascinating one-page case study described a 52-year-old woman who reported life-long facial distortions, often perceiving faces as “dragon-like.” After prolonged viewing (i.e. several minutes), faces reportedly “turned black, grew long, pointy ears and a protruding snout, and displayed reptiloid skin” ^[12]^. Neuroimaging revealed minor white-matter irregularities near the lentiform nucleus and semioval center but no other structural or functional abnormalities.

Although the case by Blom et al. (2014) ^[12]^ remains the sole published instance of non-acquired PMO, our laboratory is in contact with ten individuals who appear to have lifelong distortions. All report experiencing distortions for as long as they can remember, and several have a family history of facial distortions (see section 4.1). We refer to these cases as early-emerging PMO (EE-PMO). Their existence suggests that EE-PMO may be more prevalent than the literature indicates, much like developmental prosopagnosia was before a series of studies were published in the early 21st century ^[10]^. Like the case described below, it is often challenging to determine whether deficits in early-emerging disorders arise from early brain damage (due to an event or series of events) or developmental processes that did not operate typically.

Here, we present the first comprehensive report of EE-PMO. Zed (stage name) has experienced persistent facial distortions and face recognition difficulties for as long as he can remember. His distortions are dynamic and vary in appearance and severity (see section 2.1 and Figure 1). To investigate Zed’s condition, we conducted extensive behavioral assessments of his general visual perception, face perception and recognition, and object and voice perception and recognition. We also examined how different stimulus types and viewing conditions influenced the intensity of his facial distortions. Finally, we employed functional magnetic resonance imaging (fMRI) and diffusion tensor imaging (DTI) to explore the neural basis of Zed’s atypical face processing.

**Figure 1.**
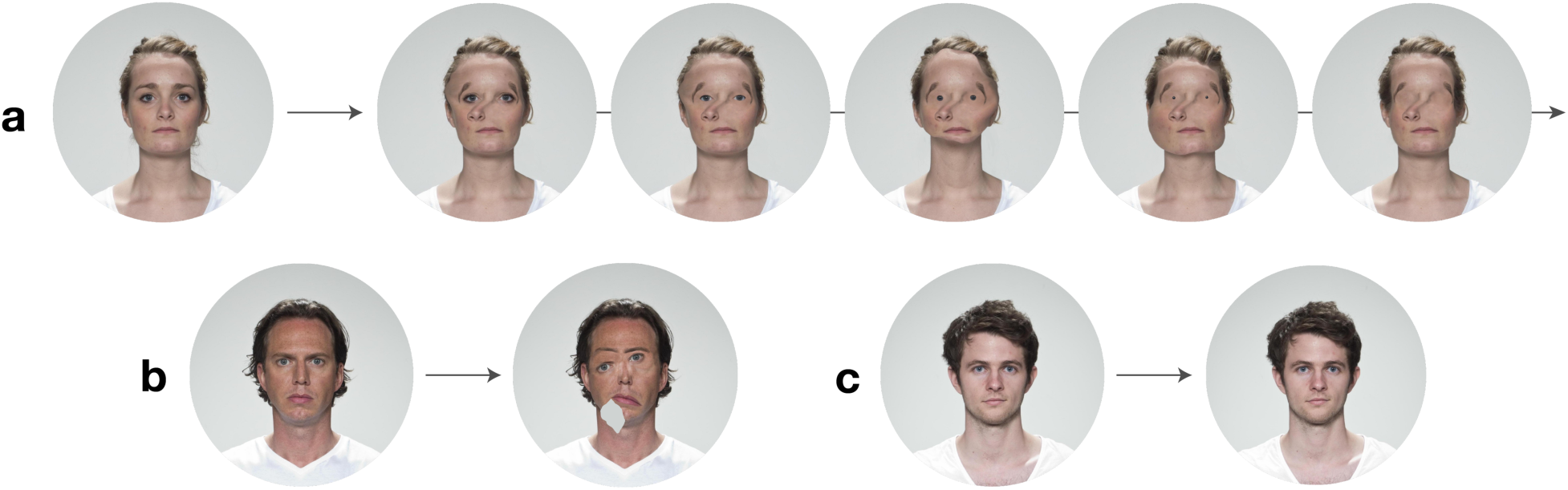
Visualizations of Zed’s face distortions. **(A)** Stills from the dynamic visualization of one of Zed’s distortions. To the left is the original face ^[36]^. Zed perceives the images on the timeline to the right morphing into one another over ∼10 seconds. When describing these distortions, he said, “the chin grows and shrinks, and then the sides of the face follow… The nose is moved to the left … The eyebrows are rotated to the sides of the eyes. Oh, and the eyes disappeared slowly, with the corners first and then the iris and then the pupil.” **(B)** A snapshot visualization of Zed’s dynamic distortions. On the left is the original face, on the right is a visualization of Zed’s distortions. Zed described this distortion in two steps. First, the “mouth droops down like it’s hinging from the left corner… and I don’t see anything in this [points to the grey area] section of the face. It’s the same color as the background.” Afterward, “the eyes are diagonal, with the left lower than the right… The eyebrows didn’t follow but I think they’re a little to the left.” **(C)** A second snapshot visualization of Zed’s dynamic distortions. On the left is the original face, on the right is a visualization of Zed’s distortions. Zed described that “the eyes are drooping on both sides to the middle of the cheeks… The nose is curved to the right … and the mouth is completely gone.”

## 2 Case Report: Zed

Zed, a teenage boy at the time of testing, is right-handed and has a history of dyslexia, aphantasia, and color blindness. His mother describes him as independent and easygoing. In all interactions with our lab, Zed has been pleasant, and he occasionally displayed his quick wit and sense of humor. He is a high school student, and enjoys music and history.

Upon entering middle school, Zed was referred to a neuropsychologist for his dyslexia. The neuropsychological evaluation indicated that Zed’s intellectual ability, theory of mind, memory, and visual-spatial ability were intact. However, he scored well below average on tests of face discrimination and recognition (5th percentile; Warrington Recognition Memory for Faces Test; Warrington, 1984) as well as long-term face memory (9th percentile; Delayed Warrington Recognition Memory for Faces Test), providing the first formal evidence of his difficulty with faces. Zed’s struggles extend beyond face recognition, though, as he also experiences facial distortions. His parent recalls that as early as kindergarten, Zed mentioned seeing “curvy noses,” “drooping eyes,” and “disappearing features.” These descriptions made it clear to her that Zed was seeing the world differently from others.

Zed is a twin and was born prematurely at 31 weeks gestational age by cesarean section, weighing 3 lbs. 5 oz. He stayed in the hospital for a month post-delivery and was treated for jaundice and provided with incubator care. After he was taken home from the hospital, Zed made good developmental progress. Apart from his birth history, Zed’s family knows of no neurological events or brain trauma that could account for his distortions, and none of Zed’s family members have experienced similar visual distortions. While all developmental milestones were attained within normal limits, Zed’s PMO may have been produced by the effects of his premature birth.

In April 2023, Zed responded to our lab’s online PMO questionnaire at https://PMO.faceblind.org/. He wrote that “sometimes faces just don’t look right I cannot describe it… it is a daily occurrence for as long as I can remember.” After receiving his response, our lab met with Zed and his mother for an online interview in which he described his experiences. He noted that his distortions are usually bilateral and tend to include facial features moving, drooping, growing, shrinking, and disappearing. Faces in his periphery often appear as blank skin and are void of any facial features. Throughout Zed’s testing, he has reported distortions on all full faces presented to him. Zed has also reported distortions in doll faces, symbolic faces, cars, animals, human bodies, and objects. Zed consistently reports dynamic distortions that change from moment to moment, and typically emerge within three seconds of viewing a face. See section 2.1 and Supplementary Video S1 for information about the visualizations of Zed’s dynamic distortions.

Unlike most PMO cases, Zed also experiences facial identity misrecognition, in which he sees incorrect identities in others ^[2]^. These misrecognitions can manifest in various ways, such as seeing familiar faces on strangers or unfamiliar faces on familiar individuals. In some cases, these misrecognitions are extreme. For instance, while in public, Zed recently approached a person whom he thought was his mother— to him, the face appeared identical to hers. Although the person’s clothes and hair were slightly different, Zed thought nothing of it. When he asked a sibling, they pointed out that the individual was, instead, a male in his 60s. Even after the correction, Zed could not see the man’s true appearance—he only saw his mother’s face. These identity misrecognitions can occur multiple times per day. He often relies on hair to recognize and remember individuals, but when contextual cues are insufficient for correction, he seeks confirmation from others before approaching someone he might know. When asked to identify the misrecognized faces, Zed reports the context in which he typically encounters the person (e.g., “they are in my orchestra”) rather than their actual name. At times, he has also perceived familiar people as strangers and has misrecognized himself in the mirror, stating it “doesn’t look like me.” Despite his distortions and misrecognitions, Zed is socially successful. He has friends at his high school, and he reports that he recognizes them via their hair, clothing, or body characteristics.

### 2.1 Distortion Visualizations

Given the inherently subjective nature of PMO, quantifying an individual’s experience is challenging. Zed’s distortions are dynamic and variable, which makes them difficult to describe to others. To address this, we created the visualizations shown in Figure 1. As with most PMO cases, Zed sees distortions regardless of the viewing medium (e.g., photographs, television, or in-person ^[14]^). Therefore, Zed cannot be certain that the visualizations accurately represent his distortions. However, he believes that these visualizations are good approximations of the distortions he perceived and may help others better understand his experience.

The preliminary visualization procedure took place in person. We selected stimuli that consistently elicited severe distortions (refer to Section 3.6.1). While looking at one image, he described each distortion he perceived, and we then manipulated a second image of the face to match his description. Throughout the in-person session, Zed provided feedback on the placement, size, and picture-plane orientation of each facial feature to achieve relatively accurate representations of each distortion. For the female face (seen in Fig 1A), which involved dynamic distortions, we followed the same procedure, with Zed providing detailed descriptions of the transitions between each distortion over time.

After the preliminary visualization procedure, each face was modified using Adobe Photoshop and After Effects. In a subsequent session, Zed reviewed the drafted visualizations and provided additional feedback. Despite perceiving additional distortions that impacted his representation of the visualizations, Zed still confidently offered constructive feedback regarding their accuracy. This feedback process continued until Zed felt that the visualizations effectively portrayed his experience. Refer to the video morph of Zed’s temporal distortions in the supplementary material under Resource A2.

## 3 Methods and Results

Following the initial screening, Zed remotely completed a background battery of standardized tests and questionnaires available online via Testable ^[15]^. The control data, Zed’s performance, and the statistical tests conducted for these tests, as well as for others mentioned above, are presented in Supplementary Table S1.

### 3.1 Low-Level Vision Testing

Zed’s low-level visual abilities are intact, as evidenced by his performance in the Hanover Early Vision Assessment ^[16]^ (HEVA) with a score of 83% (control mean = 74.6%, SD = 10.1%, t = 0.82).

During an in-person assessment, Zed correctly identified 5 out of 25 plates in the Ishihara Test ^[17]^, which detects red-green color deficiencies. His results were not specific to one particular type of color deficiency.

Zed’s impaired color perception are also evidenced by his significantly elevated total error score of 164 (control mean = 65, SD = 14.7, t = -6.62) on the Farnsworth Munsell 100-Hue Test for Color Vision ^[18,19]^, demonstrating impaired color discrimination ability.

To assess his visual imagery, Zed took the Visual Vividness Imagery Questionnaire (VVIQ). The VVIQ asks individuals to imagine an object, scene, or friend, and rate the vividness of the image on a Likert scale ^[20,21]^. Zed scored significantly below controls with a mean rating of 0.8 out of 5 (control mean = 3.35, SD = 0.93, t = -2.73), indicating that he is aphantasic and lacks low-level sensory visual imagery ^[22,23]^.

### 3.2 Face detection

Next, we evaluated Zed’s face detection, often considered the first stage of face processing ^[24,25]^. He completed two two-tone forced-choice tasks designed to measure face detection ability. Additional statistical analyses for each measure are provided in Supplementary Table S1.

Face Mooney Forced-Choice: Zed’s face detection ability was tested using the Face Mooney 3AFC, a forced-choice task in which he was presented with three two-tone images simultaneously for 3000 ms and asked to identify which image contained a face ^[26,27]^. Zed scored normally for upright (82%, control mean = 87.5%, SD = 5.65%, t = -0.95) and inverted (64%, control mean = 61.2%, SD = 10.91%, t = 0.26) faces.

Two-tone Forced-Choice: Participants viewed three black-and-white stimuli simultaneously for 500 ms and were asked to select the stimulus that contained a face ^[26,27]^. Zed scored significantly below controls on this assessment (82%, control mean = 95.8%, SD = 5.54%, t = - 2.44), which suggests he may have difficulties with face detection.

### 3.3 Face recognition

To assess Zed’s ability to recognize faces, he completed four face recognition tests. Figure 2 and Supplementary Table S1 provide additional information regarding statistical analyses, the control participants, and the control results for each of the face recognition measures.

**Figure 2.**
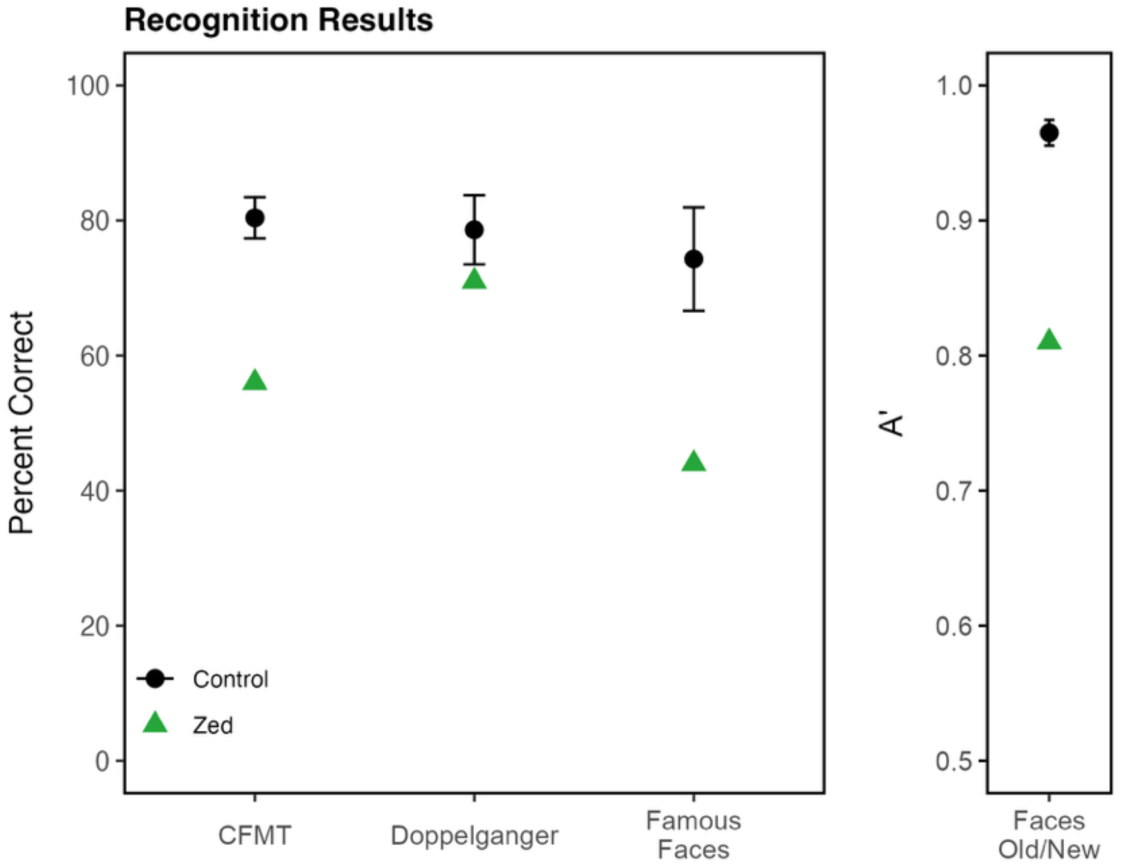
Results of the face recognition tests. Zed (green) scored significantly below controls (black) on all face recognition tests. Error bars represent ±2 standard deviations around the control mean.

#### Cambridge Face Memory Test (CFMT)

The CFMT presents a series of six unfamiliar target faces before presenting test items that consist of three faces, one of which is one of the target faces (CFMT) ^[28]^. Zed’s task was to pick out the target face. He scored significantly below the controls (56%, control mean = 80.4%, SD = 11%, t = -2.20).

#### Faces Old/New Test

Zed attempted to memorize 10 unfamiliar target faces, with each shown twice during the learning phase ^[29]^. Then, he was presented with 50 faces one at a time: 20 old faces (10 target faces each presented twice) and 30 new faces. For each face, Zed indicated if it was old or new. These old/new results were quantified using A-prime, with values ranging from chance discrimination (0.5) to perfect discrimination (1.0). Zed A-prime was 0.81, which is significantly below control scores (control mean A’ = 0.965, SD = 0.02, t = -7.53).

#### Famous Faces Test

Zed viewed 60 celebrity faces, which were closely cropped so only the face was visible ^[30]^. Of the 27 celebrities Zed was familiar with, he identified 44% of them correctly, which is two standard deviations below the control mean (control mean = 74.3%, SD = 15.1%, t = -1.94).

#### Doppelganger Test

In the Doppelganger Test, Zed was presented with the name of a famous person, followed by two simultaneously presented photos—one of the celebrity, and another of a lookalike of that celebrity ^[30]^. Zed correctly identified 71% of the 42 celebrities he was familiar with. This score was in the normal range (control mean = 78.6%, SD = 10.3%, t = -0.71).

### 3.4 Face perception

Zed’s impaired face recognition performance motivated assessment of his face perception. This testing explores whether the recognition impairments arise due to face perception or face memory deficits. Additionally, these measures will allow assessment of the extent to which Zed’s distortions impact judgments based on face perception. Refer to Fig 3 and Supplementary Table S1 for additional information about each of the face perception measures.

**Figure 3.**
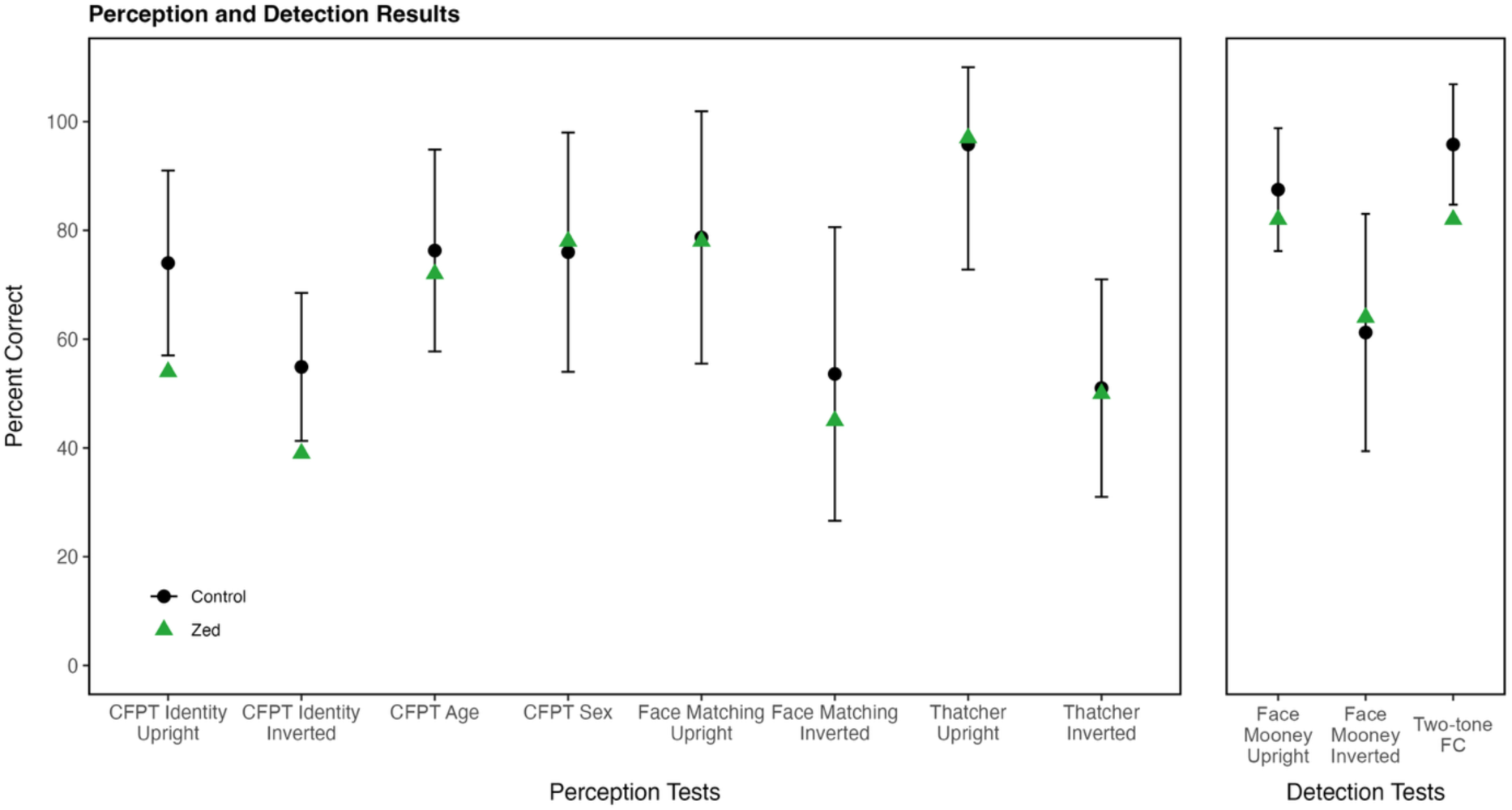
Results of face perception and detection tests. Zed’s scores (green) are compared to control means (black) across face perception and detection tests. Error bars represent ±2 standard deviations around the control mean.

#### Cambridge Face Perception Test (CFPT)

Three aspects of Zed’s face perception were evaluated using tests based on the format of the Cambridge Face Perception Test (CFPT) ^[31]^. Each assessment required Zed to sort faces by identity, age, or sex. In the CFPT-Identity, Zed attempted to arrange six facial images based on their similarity to a target face. The six images were created by morphing the target face with six unique faces in different proportions. Scores for each item were calculated by summing the deviations from the correct position for each face. For the age and sex tasks, Zed attempted to sort morphed faces according to age or sex (refer to Supplementary Figures S1 and S2 for example stimuli). Zed performed significantly below controls on the CFPT-Identity for upright (54%, control mean = 74%, SD = 8.47%, *t* = -2.27) and inverted (39%, control mean = 54.86%, SD = 6.80%, *t* = -2.27) faces. However, Zed performed normally on the CFPT-Age (72%, control mean = 76.3%, SD = 9.28%, *t* = -0.41) and CFPT-Sex (78%, control mean = 76%, SD = 11%, *t* = 0.15). These results indicate that Zed experiences difficulty with identity perception, but surprisingly, he appears to be sensitive to characteristics that indicate age or sex differences.

#### Face Matching

In this matching task, a frontal view of a target face was presented for 600ms, and then three faces were displayed from a different viewpoint for 4000ms. Participants attempted to select the target face. Zed scored normally on the face matching task (78%, control mean = 78.7%, SD = 11.6%, *t* = -0.01).

#### Thatcher Test

In this task, two normal faces and one Thatcherized face were presented, and participants were asked to select the Thatcherized face. Thatcherized faces had eyes, mouths, or both features rotated 180 degrees within the image ^[26]^. In upright faces, the inverted features are easy to detect, but when faces are inverted, the flipped features are difficult to notice ^[32]^ (see Supplementary Figure S3 for example stimuli). Zed’s scores were comparable to the controls’ scores for upright (97%, control mean = 95.8%, SD = 11.5%, *t* = 0.07) and inverted (50%, control mean = 51%, SD = 10%, *t* = -0.10) faces.

### 3.5 Car recognition, perception, and detection and voice recognition

Zed completed car recognition, perception, and detection tasks to assess whether he has difficulties with object recognition, as well as a voice recognition task to evaluate his non-visual social recognition. For further information about the voice and car measures, see Supplementary Table S1.

#### Cars Old/New Test

This test used the same procedure as the face old/new test described above ^[29]^. Zed’s A-prime was 0.85 (control mean = 0.94, SD = 0.04, *t* = -2.24), which is significantly below controls.

#### Cambridge Car Memory Test

The Cambridge Car Memory Test ^[33]^ requires participants to learn six target cars and then select them from three-car forced-choice items. Zed performed in the low-normal range (56%, control mean = 70.6%, SD = 9.93%, *t* = -1.51).

#### Car Matching

Car perception was tested using a task designed like the face matching test above ^[27]^. Zed’s score for upright car matching was normal (78%, control mean = 76.5%, SD = 10.1%, *t* = 0.10).

#### Car Mooney Forced-Choice

To test Zed’s car detection, he was presented with three two-tone images simultaneously and asked to identify which image contained a car ^[27]^. Zed’s upright score was above the control average (90%, control mean = 75.1%, SD = 11.7%, *t* = 1.24).

#### Voices Old/New Test

Zed was tested with an old/new test for voices, in which he learned the voices of six female speakers ^[34]^. After learning the voices, he was tested on his recognition of the voices and his ability to differentiate between them. Zed scored within the normal range on all tests of voice learning and recognition, with details in Supplementary Table S1.

### 3.6 Distortion Testing

#### 3.6.1 Multiview Face Distortion Assessment

Previous studies of PMO have found that distortions may be modulated by how the faces are presented ^[35]^. We created the Multiview Face Distortion Assessment (MFDA) using the Face Research Lab London Set ^[36]^ to assess the impact of five presentation variables on PMO distortions. Each participant is shown five sets of face images, each varying in one of the following dimensions: face viewpoint, picture-plane orientation, visual angle, visual field position, and facial features visible. Each set includes between 48 and 120 photographs. After viewing each face, participants rate the severity of their distortions on a Likert-type scale from 0 to 6 (0 = No distortion, 6 = Extreme distortion). Zed reported that his severity ratings typically reflected the number of distortions occurring simultaneously on a face (See section 4.2 for details).

Each condition of the MFDA shows a frontal face at the center of the participant’s fixation as a control measure. Across all frontal fixation faces, Zed’s average distortion severity score was 2.73 out of 6, establishing Zed’s baseline level of distortion severity when perceiving a face without varied viewpoints. Zed’s distortions were not altered by changes in picture-plane orientation, visual angle, or viewpoint conditions (Supplementary Table S2). Thus, the results of these conditions are not discussed further.

In the visual field position condition, Zed fixated on a central cross and was presented with peripheral faces in the four quadrants of the screen. He experienced a 1.23-point increase in average distortion intensity rating when presented with frontal faces in his periphery (Center: M = 2.92, SEM = 0.14; Periphery: M = 4.15, SEM = 0.09). The intensity ratings at all four of the peripheral positions exceeded his average distortion severity rating for faces at fixation (Figure 4; Supplementary Table S3). These results indicate that faces in Zed’s periphery are more severely distorted than those at fixation, consistent with Zed’s reports of this phenomenon in daily life described in Section 2.

**Figure 4.**
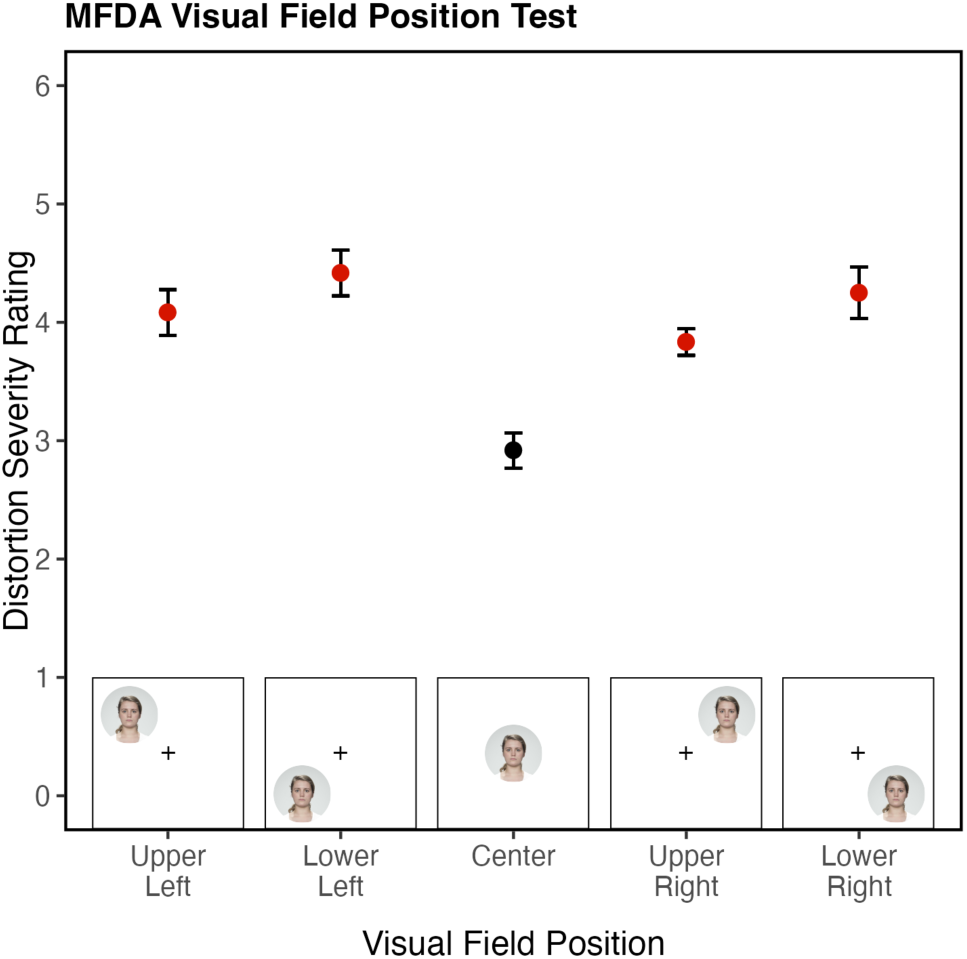
Results of the MFDA Visual Field Position Condition. Zed’s distortion severity varies depending on the position of a face in his visual field. Peripheral faces (red) elicited more severe distortions than central faces (black). Error bars represent ±1 SEM. Ratings were made on a 0–6 Likert scale.

In the visible facial features condition, Zed viewed different combinations of facial features at fixation. He experienced a decrease in distortion intensity when viewing individual features (nose, left eye, right eye, mouth; Fig 5; Supplementary Tables S3 & S4), largely due to trials in which he reported no distortions. Once two or more features were presented, Zed’s average distortion intensity was comparable to that of the full face.

**Figure 5.**
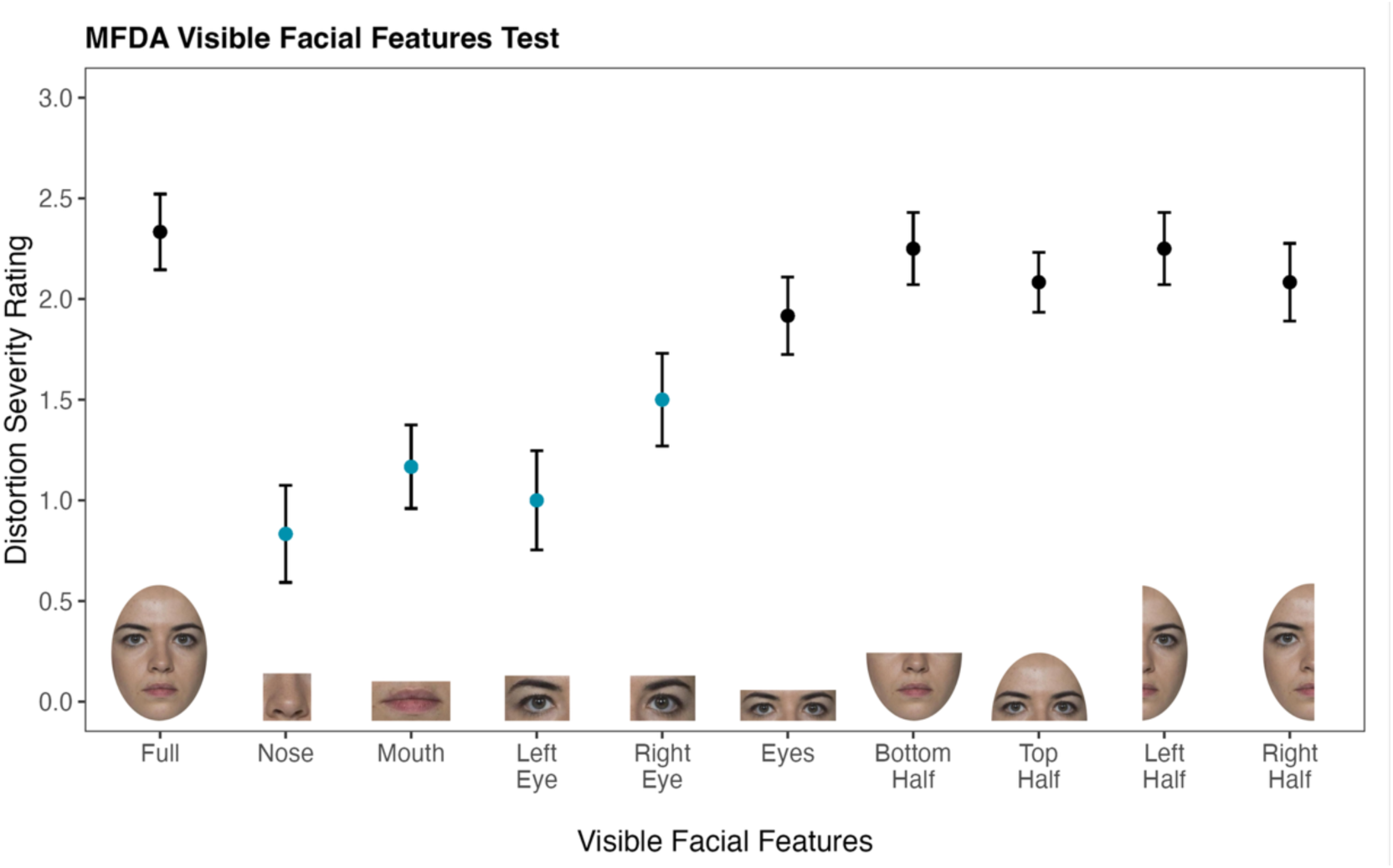
Results of the MFDA Visible Facial Features Condition. Zed’s distortion severity ratings are influenced by combinations of visible features. The y-axis (0–3) reflects the range of Zed’s ratings in this condition. Error bars represent ±1 SEM. Individual features (blue; nose, left eye, right eye, mouth) elicited less severe distortions than combined features or whole faces (black). Ratings were made on a 0–6 Likert scale.

#### 3.6.2 Familiarity Test

The familiarity test sought to determine the impact of personal familiarity on Zed’s distortion intensity, as some individuals with PMO experience distortions influenced by face familiarity ^[37]^. This test was also motivated by Zed’s frequent misidentifications of individuals in everyday life.

The familiarity test had 26 faces in total, 14 personally familiar and 12 unfamiliar. Each image was cropped around the face to exclude all hair and clothing. Each face was displayed four times, so 56 familiar and 48 unfamiliar stimuli were presented. The unfamiliar images, taken from face databases ^[38–40]^, were slightly filtered to match the unprofessional photographic appearance of the familiar images. After viewing a face for 10 seconds, Zed rated his distortion severity on the 0-6 Likert scale described above and then reported if the individual was familiar or unfamiliar. If the individual was familiar to Zed, he was prompted to name them.

Zed’s ability to discriminate between familiar and unfamiliar faces was assessed using A-Prime (chance discrimination = 0.5; perfect discrimination = 1.0). Given that a typical individual can accurately recognize personally familiar faces with ease, Zed’s A-prime score of 0.72 indicates difficulty discriminating between personally familiar and unfamiliar faces.

Zed stated that 88 of the 104 stimuli were familiar to him. His responses produced a hit rate of 92.7% (familiar face seen as familiar) and a false alarm rate of 75% (unfamiliar face seen as familiar). Of Zed’s false alarms, only 55% were supported by a specific name, while the remainder included only contextual information (i.e. “someone from school”). 36% of his false alarms involved someone he is personally familiar with. While false alarms were Zed’s primary deficit, Zed also had difficulty identifying the personally familiar individuals. Of the 56 personally familiar individuals, Zed identified only 79% correctly when asked to name them.

Zed’s distortion severity rating did not differ by familiarity. Unfamiliar faces averaged 4.08 (SD = 0.87), while familiar faces averaged 3.91 (SD = 0.67). Additionally, his recognition ability did not differ by distortion severity rating (Correctly Recognized: M = 4.14, SD = 0.62; Incorrectly Recognized: M = 3.83, SD = 0.77). Thus, Zed shows no evidence of familiarity effects on distortion severity.

### 3.7 Neuroimaging

#### 3.7.1 MRI / fMRI Methods

##### Participants

Zed and 20 neurotypical participants (14 females, mean age = 38.18, SD = 33.86) participated in this portion of the study. Written informed consent was obtained from all participants in accordance with the Declaration of Helsinki and a protocol approved by the Dartmouth College Committee for the Protection of Human Subjects.

##### Visual Stimuli and Task

To measure face selectivity, a category localizer was run that presented six classes of stimuli: faces, bodies, objects, phase-scrambled objects, natural scenes, and words. Because of Zed’s deficits with faces, our analysis focuses solely on face selectivity. Stimuli were dynamic video clips featuring various visual elements in motion, filmed against a uniform black backdrop to maximize contrast and visibility. Video clips subtended 26 x 15 degrees of visual angle in width and height. Participants viewed four runs, which each lasted 8 minutes and 22 seconds. Within each run, stimuli were grouped into 14-second single-category blocks containing five videos per block. Blocks of each category were displayed four times in each run in quasi-random order, twice in color and twice in grayscale. The trial order was the same for all participants. Throughout the experiment, participants were allowed to freely move their eyes and instructed to identify back-to-back repeats with a button press (1-back task). Each of the four runs contained unique stimuli.

##### MRI Data Acquisition

Zed and the control participants were scanned at the Dartmouth Brain Imaging Center at Dartmouth College on a 3 Tesla Siemens Prisma MRI scanner (Siemens Medical Solutions) equipped with a 32-channel receive-only phased array head coil. High-resolution whole-brain anatomical images were collected using a T1-weighted magnetization prepared rapid acquisition gradient echo (MPRAGE) sequence (208 sagittal slices; 1mm isovoxel resolution, field of view = 256mm; TR = 2,300 ms; TE = 2.03 ms; TI = 900 ms; flip angle = 9 degrees; FOV = 256 x 256 mm; bandwidth = 240 Hz/px).

Functional scans were acquired using a T2*-weighted gradient recalled echoplanar imaging multiband pulse sequence (cmrrmbep2bold) from the University of Minnesota’s Center for Magnetic Resonance Research (69 slices; nominal spatial resolution 2mm x 2mm x 2mm; 106×106 matrix size; 2 mm slice thickness, no-gap; field of view = 212mm; phase partial Fourier scheme of 6/8; TR = 2,000 ms; TE = 30 ms; flip angle = 79 degrees; bandwidth = 1814 Hz/Px; echo-spacing = 0.66 ms; excite pulse duration = 8,200 microseconds; multi-band factor = 3; phase encoding direction = AP; fat saturation on; advanced shim mode on; no iPat). Slices were oriented approximately in plane with the calcarine sulcus to ensure coverage of most, and often all, of the temporal, parietal, and occipital lobes. At the beginning of each session, a pair of EPI images with phase encoding directions of opposite polarity in the anterior-to-posterior plane were collected for post-hoc correction of EPI spatial distortion.

##### Stimulus Display and Scanner Peripherals

Stimuli were presented using a Panasonic PT-DW750BU 7000-Lumen WXGA DLP Projector and an SR Research in-bore back projection screen positioned just inside the head end of the magnet. Participants viewed the screen via a mirror mounted atop the head coil. The viewing distance was 127 mm from the eye to the mirror + 1,010 mm from the mirror to the screen = 1,137 mm total. The maximum square extent of the image projected on the screen was 297 mm x 297 mm. This resulted in a maximum possible visual angle of 14.97 degrees.

A Lenovo Thinkpad T480s computer running Linux Ubuntu 20.04.6 (Focal Fossa) controlled stimulus presentation and recorded button presses using Matlab (R2021B) code and PsychToolbox (lookup version) extensions. Behavioral responses from scanning sessions were collected using a Current Designs two-button fiber-optic handheld response pad, connected to a Current Designs 932 interface.

##### Preprocessing of fMRI data

All DICOM images were converted to NIfTI format using dcm2niix version 1.0.2018.11.25 (Li2016_JNM). From each participant’s T1-weighted volume, cortical surface meshes of the pial-gray matter boundaries and gray matter-white matter boundaries were reconstructed using Freesurfer’s recon-all program ^[41]^.

Subsequent pre-processing was performed using AFNI version 23.3.12 Septimius Severus ^[42]^ and AFNI’s afni_proc.py. Slice timing differences were corrected, and non-linear geometric distortion correction was applied using separately acquired volumes where the phase-encoding direction was reversed (blip-up/blip-down strategy). EPI-anatomical alignment was carried using the lpc+ZZ cost function while checking for possible left-right flips ^[43]^. Motion correction was applied, and the parameter estimates were stored for later use as nuisance regressors.

Timepoints were censored when either the Euclidean norm exceeded 0.3 mm or when the outlier fraction exceeded 5%. Data were then iteratively smoothed to achieve a uniform smoothness of 4mm FWHM (twice the EPI voxel dimension). The volumetric data was then projected onto the corresponding hemispheres of the high-density SUMA standard meshes (created from the Freesurfer output with AFNI function @SUMA_Make_Spec_FS). Finally, each time series was scaled to units of local BOLD percent signal change. All pre-processing results were carefully visually inspected using the QC HTML reports generated by afni_proc.py ^[43]^.

##### Face-Selectivity Analysis

To estimate the face-selectivity of 14 face-selective areas in Zed and each control, we employed the following cross-validated procedure ^[44,45]^. The four runs of the dynamic functional localizer were split into selection and test runs in a “leave-one-run-out” method chosen to avoid issues of circularity. In each cross-validated fold, three runs were used for voxel selection, and the remaining run was used to compare the response to faces and objects. To select face-selective voxels, we computed the statistical contrast between blocks of faces and blocks of objects and windowed the results by seven different region of interest (ROI) masks in each hemisphere. Face-selective ROI masks included three ventral temporal regions (the occipital face area (OFA), the posterior fusiform face area (pFUS), and the medial fusiform face area (mFUS), two lateral regions (anterior superior temporal sulcus face area (aSTS), and the posterior superior temporal sulcus face area (pSTS), and two regions of the extended face processing network (the anterior temporal lobe face area (ATL), and the inferior frontal gyrus face area (IFG)). Masks were defined in group space (fsaverage6) based on face localizers collected from a separate group of pilot participants. Within each mask, we ranked voxels by their *t*-value for the faces-vs-objects contrast across the three selection runs and selected the top 10% of voxels with the highest values. Within this set of selected voxels, we then measured the difference between beta coefficients (scaled to percent signal change) for face and object blocks in the left-out run. The final measure of category selectivity was the average of the measurements across all four cross-validated folds. Bayesian tests of deficits in the presence of covariates ^[46]^ were used to test for differences between Zed’s category selectivity and that of the 20 controls, controlling for variation related to age.

#### 3.7.2 fMRI Results

Figure 6 displays a whole-brain GLM contrasting blocks of faces versus blocks of objects for Zed. We found face-selective clusters of activity bilaterally in the inferior occipital gyrus, fusiform gyrus, superior temporal sulcus, anterior temporal lobe, and inferior frontal gyrus, with anatomical locations that are well aligned with previously studied nodes of the face-selective network ^[47,48]^.

**Figure 6.**
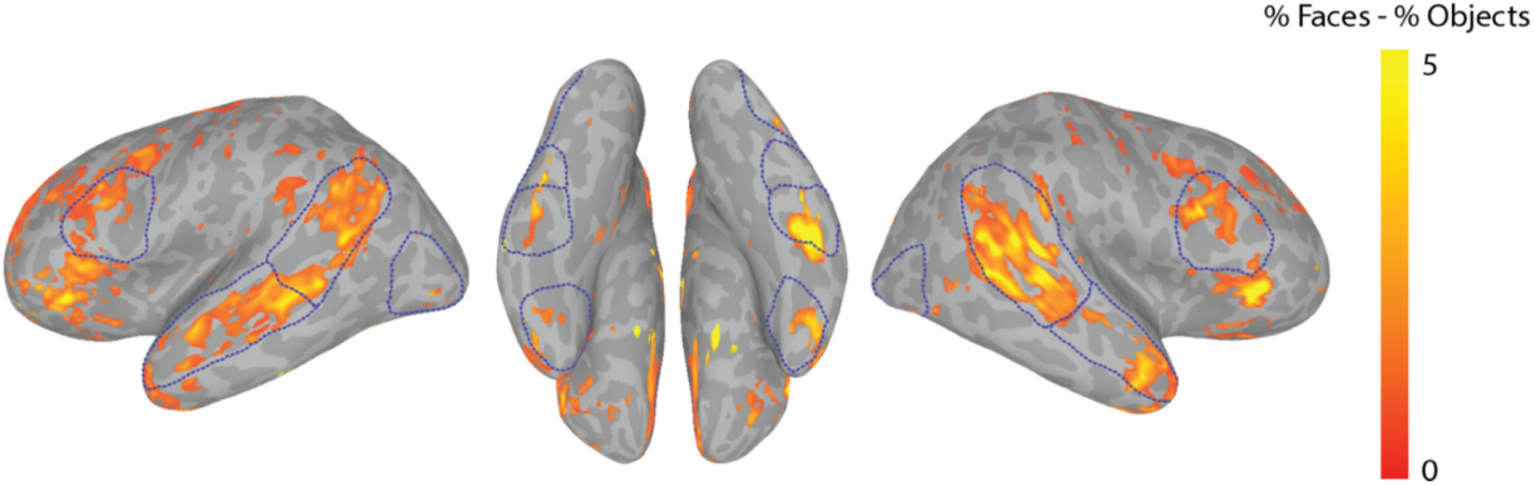
Zed’s whole-brain GLM analysis with colors indicating the difference in percent signal change for faces minus the percent signal change for objects following thresholding (*t* > 3.3, *p* < 0.001, uncorrected). Group-defined masks used for quantifying ROI-based face selectivity are outlined in blue. Group masks were drawn on the *fsaverage6* template and displayed on Zed’s standardized meshes following alignment based on sulci and gyri.

To quantitatively compare face selectivity in Zed and the neurotypical controls, we used the variable window method ^[45]^ to examine the mean percent signal change to faces minus objects within the group-defined region of interest (ROI) masks ^[44]^ (Figure 7). Separate masks were created for the posterior and anterior aspects of the fusiform gyrus (pFUS/FFA-1 and mFUS/FFA-2, respectively) using the mid-fusiform sulcus as the separating border ^[49]^. Bayesian tests for the presence of a deficit between Zed and the control group were conducted within each ROI, using age as a covariate of no interest ^[46]^ (face selectivity as a function of age is plotted in Supplementary Figure S4). Compared to controls, Zed showed significant reductions in face selectivity in the right OFA (*Z_ccc_=* -2.01, *p* = 0.04) and right pFUS (*Z_ccc_ = -2.82*, *p* = 0.01). The reduction in face selectivity was largest in right pFUS, with an estimated 99.02% of the population having greater face selectivity than Zed (CI: 93.89% to 99.995%). Inspection of the beta coefficients for individual categories (see Supplementary Fig S5) revealed Zed had similar activation for both faces and objects in right pFUS, so his decreased face selectivity resulted from a reduced response to faces rather than an elevated response to objects. Descriptive statistics, effect sizes, and interval estimates for all ROIs are presented in Table 1.

**Figure 7.**
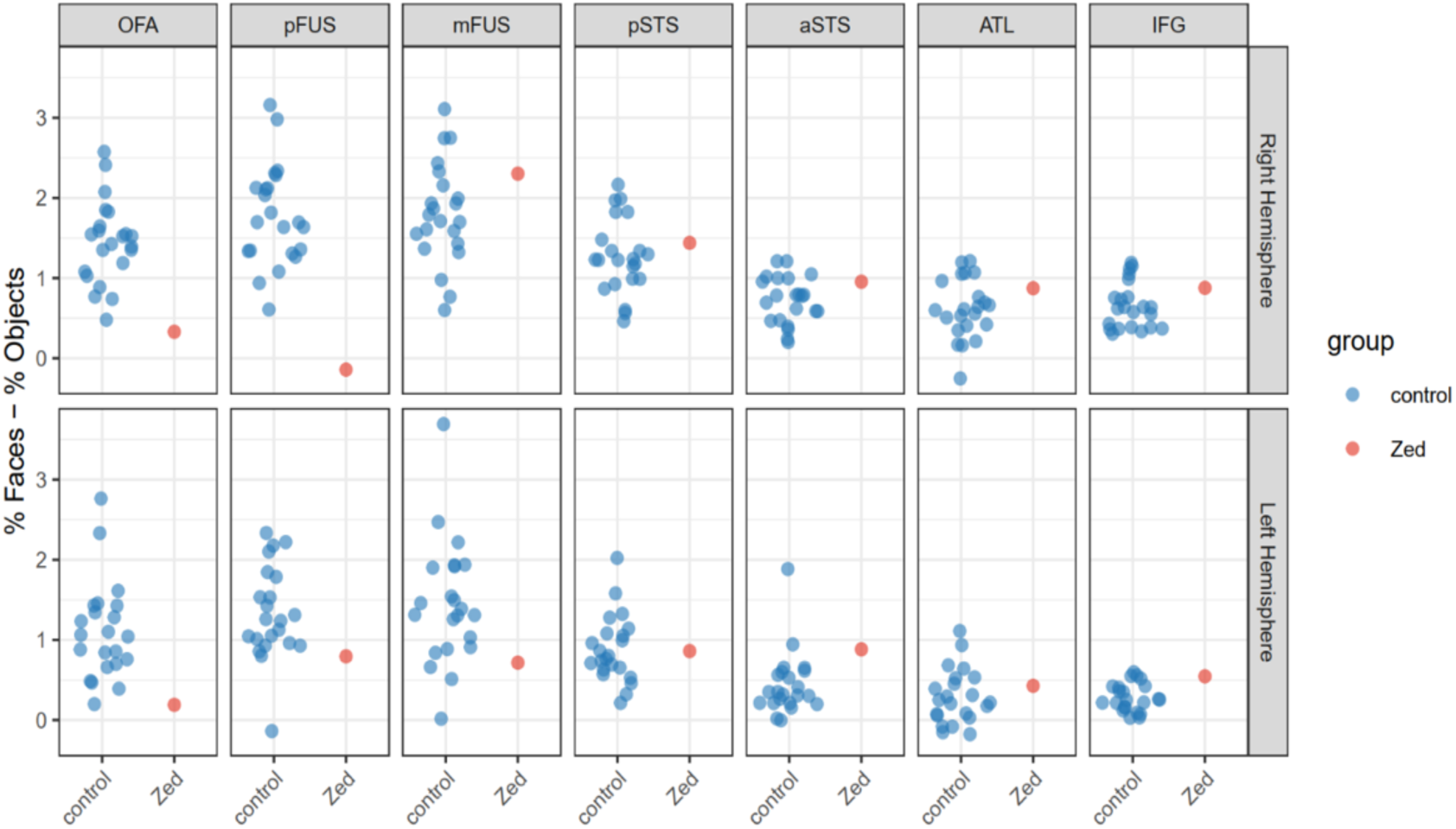
Face selectivity in Zed and Controls. Face selectivity expressed as percent signal change to faces minus percent signal change to objects for Zed and each control by ROI and hemisphere.

**Table 1.** ROI-based Bayesian tests of deficit comparing Zed to neurotypical controls. Abbreviated columns are defined as follows: Zed’s mean delta % signal change (faces vs. objects), control mean delta % signal change (faces vs. objects), control standard deviation (SD) of control delta % signal change (faces vs. objects). *Z*_ccc_ = case–control effect size ctonrolling for covariate of age. P-value is two-tailed.

| Table 1. ROI-based Bayesian tests of deficit comparing Zed to neurotypical controls. |  |  |  |  |  |  |  |  |
| --- | --- | --- | --- | --- | --- | --- | --- | --- |
| ROI | Hemisphere | Zed Mean $\Delta\%$ Signal Change | Control Mean $\Delta\%$ Signal Change | Control SD $\Delta\%$ Signal Change | $Z_{ccc}$ (95% CI) | df | p-value | Estimated % of Population Below Zed (95% CI) |
| OFA | RH | 0.33 | 1.446 | 0.516 | -2.013 (-2.933, -0.915) | 20 | **0.043** | 4.295 (0.168, 18.017) |
| OFA | LH | 0.191 | 1.106 | 0.607 | -1.494 (-2.386, -0.503) | 20 | 0.098 | 9.827 (0.851, 30.740) |
| pFUS | RH | -0.142 | 1.781 | 0.628 | -2.817 (-3.907, -1.546) | 20 | **0.01** | 0.976 (0.005, 6.110) |
| pFUS | LH | 0.795 | 1.334 | 0.582 | -0.525 (-1.362, 0.308) | 20 | 0.32 | 32.008 (8.656, 62.081) |
| mFUS | RH | 2.303 | 1.803 | 0.628 | 1.052 (0.158, 1.888) | 20 | 0.823 | 82.349 (56.257, 97.049) |
| mFUS | LH | 0.716 | 1.455 | 0.768 | -0.771 (-1.6, 0.094) | 20 | 0.247 | 24.697 (5.484, 53.749) |
| pSTS | RH | 1.439 | 1.267 | 0.465 | 0.66 (-0.213, 1.471) | 20 | 0.72 | 72.002 (41.562, 92.933) |
| pSTS | LH | 0.861 | 0.881 | 0.418 | 0.206 (-0.608, 1.034) | 20 | 0.574 | 57.373 (27.159, 84.935) |
| aSTS | RH | 0.955 | 0.727 | 0.294 | 1.972 (0.9, 2.905) | 20 | 0.955 | 95.474 (81.586, 99.816) |
| aSTS | LH | 0.883 | 0.444 | 0.395 | 1.915 (0.872, 2.873) | 20 | 0.95 | 95.045 (80.835, 99.796) |
| ATL | RH | 0.873 | 0.618 | 0.374 | 1.073 (0.163, 1.939) | 20 | 0.828 | 82.780 (56.455, 97.376) |
| ATL | LH | 0.428 | 0.295 | 0.341 | 0.799 (-0.074, 1.619) | 20 | 0.76 | 76.010 (47.036, 94.722) |
| IFG | RH | 0.877 | 0.652 | 0.287 | 0.463 (-0.392, 1.285) | 20 | 0.658 | 65.805 (34.770, 90.053) |
| IFG | LH | 0.546 | 0.285 | 0.175 | 1.331 (0.389, 2.201) | 20 | 0.879 | 87.860 (65.126, 98.612) |

#### 3.7.3 DTI Methods

##### Data Acquisition and Preprocessing

Diffusion-weighted imaging (DWI) data were acquired from Zed and 78 healthy controls using a Siemens Magnetom Prisma 3T scanner with a 32-channel head coil (Control dataset from ^[50]^). Controls averaged 25.03 years of age (SD=5.25), and ranged in age from 18 to 42. Forty-five were women, 31 were men, and two selected other gender ^[50]^. Technical difficulties with the anterior section of the head coil during Zed’s scan resulted in reduced signal-to-noise ratio (SNR) in frontal brain regions. To minimize the risk of contamination from low-quality data, we excluded the anterior 4 cm of the brain from analysis (corresponding to ∼22% of total brain length in MNI space, which spans 18 cm). As shown in the results, this exclusion did not affect the comparison between Zed’s results and those of the controls in the remaining ROIs.

The DWI protocol employed a multi-shell acquisition scheme with b-values of 0, 500, 1000, 2000, and 3000 s/mm², comprising 96 non-zero diffusion directions distributed as 6, 15, 15, and 60 directions across the respective shells. Data were acquired with 1.7 mm³ isotropic voxels and preprocessed using MRtrix3, ANTs, and FSL. Standard preprocessing steps included quality control assessment, denoising, Gibbs ringing artifact removal, motion and distortion correction using top-up, and bias field correction. The preprocessed data were regridded to 1 mm³ isotropic resolution for analysis.

##### Diffusion Model Fitting, ROI Analysis, and Statistical Comparisons

Diffusion tensor imaging (DTI) parameters were calculated using the b = 1000 s/mm² shell, generating maps of fractional anisotropy (FA), mean diffusivity (MD), axial diffusivity (AD), and radial diffusivity (RD1 and RD2). Following model fitting, all parametric maps were transformed to MNI standard space. Fifty-two white matter ROIs were defined using the Johns Hopkins University white matter atlas ^[51,52]^, which was implemented in FSL. Mean values for each diffusion metric were then extracted for all participants.

Modified t-tests designed for single-case comparisons assessed whether Zed’s values for each metric within each ROI differed significantly from the control group mean^[46,50,53]^. To address multiple comparison issues arising from testing five metrics per ROI across 52 ROIs (260 comparisons), the false discovery rate (FDR) was controlled using the Benjamini-Hochberg procedure ^[54]^, with statistical significance defined as p-FDR < 0.05. Results were interpreted in the context of white matter microstructural integrity to identify regions where Zed’s brain structure deviated from that of the controls.

#### 3.7.4 DTI Results

The DTI analysis involved five measures of white matter integrity (FA, MD, AD, RD1, RD2) for each of the 52 ROIs we analyzed. In 49 ROIs, Zed’s values across all five DTI measures did not differ significantly from those of the controls (see Supplementary Table S5). The predominance of normal values in the ROIs indicates that the head coil malfunction had minimal impact on the analyses. In contrast, significant differences in at least one measure were observed in three ROIs (Supplementary Table S5). The largest deviations were observed in the left inferior fronto-occipital fasciculus (IFOF; Figures 8 & 9; Supplementary Table S5), with differences in FA, MD, RD1, and RD2 consistent with reduced white matter integrity within this ROI. Additional significant differences were found for one measure (FA) in the right anterior coronal radiata and one measure (MD) in a combined ROI consisting of the crus of the left fornix and left stria terminalis (see Supplementary Table S5).

**Figure 8.**
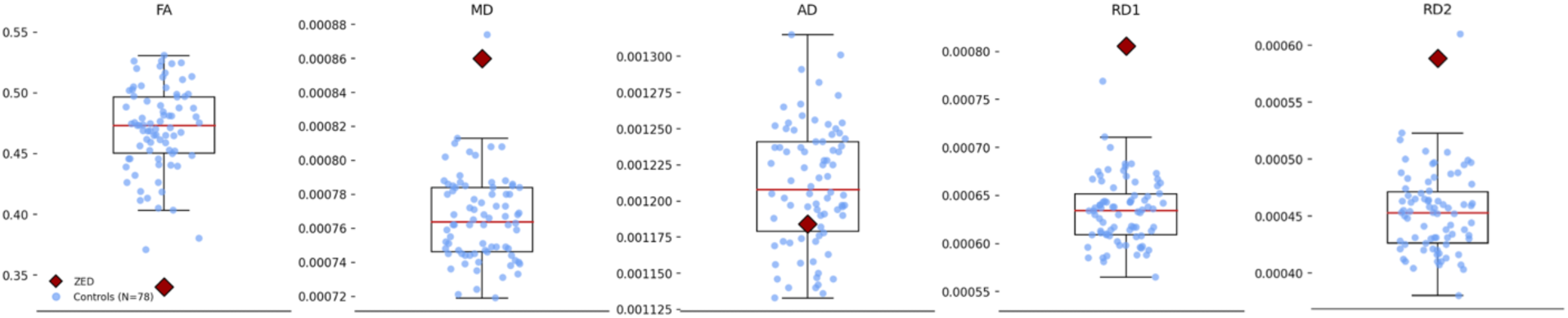
White matter abnormalities in Zed’s left inferior fronto-occipital fasciculus. Four significant differences were found in the measurements of the left inferior fronto-occipital fasciculus.

**Figure 9.**
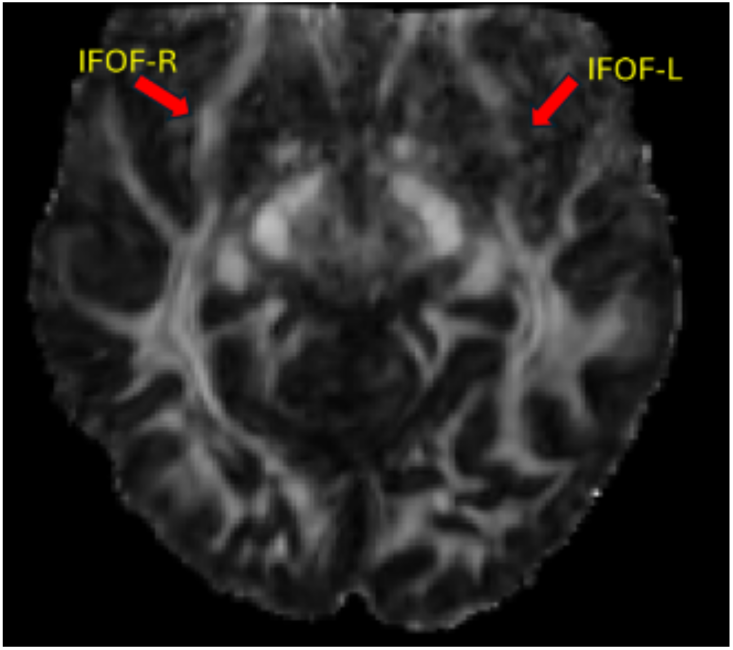
Reduced FA in Zed’s Left IFOF. The FA value for Zed’s left IFOF (IFOF-L) is significantly below that of the controls. Visual comparison of Zed’s left IFOF and right IFOF (IFOF-R) makes this difference evident.

Additional significant differences were found for two measures in both the left and right anterior coronal radiata, along with one measure in the left uncinate fasciculus, the left external capsule, and the right external capsule (see Supplementary Table S5).

## 4 Discussion

In this case study, we provide a comprehensive analysis of co-occurring PMO and prosopagnosia by assessing Zed’s distortion severity, neuropsychological profile, face-selective areas, and white-matter characteristics. Zed experiences some of the most intense distortions in the literature, and he sees them on every face he encounters. Notably, despite his pronounced facial distortions, Zed’s perceptual judgments of faces remained intact, highlighting a dissociation that warrants further investigation. Neuroimaging revealed Zed’s facial distortions are accompanied by both abnormal white matter and reduced face-selectivity in the right OFA and the right posterior FFA. Below, we discuss Zed’s results, their implications, and some of the complications in interpreting findings from participants who experience facial distortions.

### 4.1 Early-emerging prosopometamorphopsia

Zed reports lifelong facial distortions, recognition difficulties, and misidentifications. While his family is unaware of neurological events that might have caused brain damage, Zed’s distortions may stem from early brain damage resulting from his premature birth rather than atypical developmental processes. Multiple studies show that preterm individuals exhibit elevated rates of white matter abnormalities, and some of these studies suggest that the IFOF may be particularly vulnerable ^[55–60]^. As a result, given the uncertainty about the cause of Zed’s neural abnormalities, we are referring to Zed’s case as early-emerging PMO (EE-PMO) rather than early-onset acquired PMO or developmental PMO.

Although Zed and the case reported by Blom et al. (2014) ^[12]^ are the only published accounts to explicitly describe lifelong symptoms, our lab has heard from other EE-PMO cases, and we suspect that such cases are more common than the literature suggests. For instance, Ezri, a woman in her early thirties, experiences bilateral PMO that alters the shape and texture of faces ^[61]^. She does not recall a time without these distortions. When Ezri shared her experiences with her mother, her mother recalled five or six instances in which she also saw salient facial distortions. Similarly, another woman we are in touch with has reported lifelong facial and body distortions. When she discussed her symptoms with her three teenage daughters, one described episodes in which her arms appeared unusually long, while another reported seeing distorted faces. These family histories suggest a potential genetic contribution to PMO. No such familial pattern, however, is present in Zed’s case, as none of his family members report visual distortions.

In cases like Zed’s in which early-emerging distortions are dynamic and varied, it seems likely that individuals will recognize that their perceptual experiences differ from those of others.

However, if early-emerging distortions are static and consistent across viewpoints and different faces as has been observed in acquired PMO ^[62]^, individuals may not recognize their perceptions as distorted. Instead, we suspect that they are likely to interpret the altered features as accurate representations of others’ faces, and they may never realize that their perception of faces is severely distorted.

### 4.2 Identity misrecognition

In addition to experiencing distortions, Zed frequently misrecognizes identities, identifying strangers as individuals he knows. After a misrecognition during an assessment, he noted, “I did not recognize the face even after you revealed the name. I still think it’s the person I said it was.” His misrecognitions can span differences in age, sex, and race. For instance, on the Familiarity Test, he confidently identified a 70-year-old man as a 20-year-old woman. In another case, described by his mother, he approached a white man assuming he was black. While prosopagnosia may contribute to these errors, the marked differences between the actual face and the face he perceives, as well as the frequency with which he mistakenly sees familiar faces, suggest that these faces are being retrieved from memory during misrecognitions.

While no prior reports document frequent misrecognitions across age, sex, or race to the extent observed in Zed, several studies describe similar identity misrecognitions. Following a right middle cerebral artery cerebral infarction, one man reported repeated misidentifications of unfamiliar individuals as family or friends, despite performing relatively well on sex, age, and face matching tasks — similar to Zed ^[63]^. Unlike Zed, however, this individual also struggled to recognize familiar voices. In a more extreme case, a prosopagnosic woman reported that, after briefly looking at a poodle on the bus, all of the passengers’ faces had transformed into that of the poodle—a distortion that persisted for 30 minutes ^[64]^. Intracranial stimulation studies have also elicited phenomena resembling Zed’s distortions and misrecognitions, with stimulation of the right fusiform face area (FFA) inducing both facial distortions and feelings of familiarity. In one case, a patient reported facial drooping, and reported that the face “almost looks like somebody I’ve seen before” ^[6]^. Similarly, another patient described seeing the eyes, mouth, and mole of a previously presented face superimposed atop a new face following right FFA stimulation ^[65]^. In a separate trial, the patient reported that the face had “eyes and ears which were not theirs” but could not match familiar identities to these features. Notably, Zed’s right posterior FFA also exhibited marked functional anomalies, suggesting that these irregularities may underlie his distortion and misrecognition experiences.

### 4.3 PMO and face perception

Results from Zed’s behavioral and neuropsychological testing are consistent with both PMO and prosopagnosia. Although PMO is more frequently reported in individuals with prosopagnosia ^[2]^, face recognition is usually normal in PMO ^[2,12]^. Even in cases of bilateral PMO with pronounced facial distortions, recognition abilities often appear to remain intact ^[1,12,66]^. Because PMO is defined as a disorder of face perception, we evaluated Zed’s performance on face perception and detection tasks. Surprisingly, despite his pronounced facial distortions, Zed performed within the normal range on four out of five face perception measures and showed difficulty only with the CFPT-Identity, which required sorting based on facial identity. Zed’s normal performance on the CFPT-Age and CFPT-Sex was particularly unexpected. Good performance on these tests requires extended, close scrutiny of the subtle age- or sex-related differences between faces and a level of perceptual precision that we anticipated would be disrupted by his distortions.

It’s not clear from the current results how Zed is able to perform normally on fine-grained facial judgments despite his strong distortions. As noted earlier, his distortions typically emerge within three seconds of viewing a face, so it is plausible that he completed perceptual judgments before the distortions fully developed. Alternatively, these findings may point to a dissociation between perceptual experience and perceptual judgment about faces. Like Goodale and Milner’s distinction between visual perception and visual control of action ^[67]^, Zed’s distortions may arise from a system supporting conscious face perception, while his judgments may rely on a separate intact system. However, while we find this an intriguing possibility, far more evidence about the relationship between awareness of faces and judgments about faces is needed before confident conclusions can be drawn.

### 4.4 Neural correlates of Zed’s face distortions and prosopagnosia

To investigate the neural correlates of Zed’s distortions, we carried out functional and structural imaging with Zed. A localizer study assessing face-selective areas revealed that Zed exhibited notable reductions in several regions. Whole-brain analyses revealed that although Zed recruited canonical face-selective regions—such as bilateral inferior occipital gyrus (OFA), fusiform gyrus (FFA), superior temporal sulcus (STS), anterior temporal lobe, and inferior frontal gyrus—, but quantitative comparison to results from neurotypical controls revealed significantly reduced face selectivity in key nodes of the network. Specifically, Zed showed reductions in face selectivity in the right OFA and a pronounced reduction in right pFUS. The right pFUS reduction was especially pronounced. Inspection of Zed’s beta coefficients to faces and objects shows that his reduced face selectivity resulted from a weak response to faces rather than an unusually strong response to objects. Thus, despite the preserved presence of face-selective regions in both hemispheres, the functional face-selectivity—particularly in the right pFUS—was atypical.

The abnormalities observed in Zed’s right posterior face regions are notable given evidence that intracranial stimulation of these face-selective areas can alter conscious perception of faces.

Direct intracranial stimulation of the right OFA and right pFUS has repeatedly distorted the appearance of faces ^[6,7,65,68–72]^. In contrast, stimulation of face-selective regions in the left hemisphere only rarely alters conscious face perception ^[7]^, and stimulation of anterior face regions in the right hemisphere disrupts the ability to recognize facial identity without affecting facial perception ^[5,73]^. These intracranial findings suggest that right OFA and right FFA play particularly critical roles in the distortions observed in PMO cases, consistent with Zed’s results.

In addition to functional abnormalities, Zed also exhibited white-matter deficits in three regions of interest (ROI). The ROI with the clearest reductions in white matter integrity was the left fronto-occipital fasciculus. Two other ROIs, the right anterior coronal radiata and a combined ROI consisting of the crus of the left fornix and left stria terminalis, also suggested reductions in white matter integrity, but given their functional roles, they appear unlikely to contribute to Zed’s distortions. On the other hand, deficits in the left IFOF could be linked to visual distortions. The IFOF is a ventral white matter tract projecting between the occipital and frontal lobes ^[74]^ and is a crucial contributor to the language network and semantic speech processing ^[75,76]^. Subcortical stimulation of the dorsal IFOF has resulted in alexia (pure word blindness) and agraphia (loss of writing ability) ^[76]^. In addition, some findings suggest it may be involved in object recognition ^[77]^ and face processing ^[78]^. In adolescents, Taddei et al. (2012) found significant correlations between IFOF FA values and normalized N400 amplitude in response to angry faces ^[78]^.

Additionally, Philippi et al. (2009) found that damage to the right IFOF predicted facial emotion recognition impairments for negative emotions such as sadness, anger, and fear ^[79]^. That study also presented a subject with a pure white matter lesion of the right IFOF who possessed a specific facial expression recognition deficit. Due to the IFOF’s multifaceted role in semantic language processing *and* face processing, it is not possible to confidently connect Zed’s IFOF findings with a single behavioral output, but it is likely that Zed’s abnormal left IFOF is implicated in his comorbid dyslexia, face recognition difficulties, and face distortions.

## 5 Data Availability

All neuropsychological, behavioral, and control data (collected on Prolific) are available on the Open Science Framework (OSF): https://osf.io/d5ej9/?view_only=328304d992514be4b200137101a5347c. Zed’s DTI and fMRI data are available on OpenNeuro: https://openneuro.org/datasets/ds006471. Control DTI are available on OpenNeuro under sub-XXXX/ses-01/anat/dwi: https://openneuro.org/datasets/ds005256/versions/1.1.0. Control fMRI data are available on OpenNeuro: https://openneuro.org/datasets/ds006472.

## 7 Author Contributions Statement

S.F., A.M., and B.D. conceived the study. S.F., A.M., D.S., and B.D. designed the paradigms. S.F. performed data collection. H.J. and D.S. collected and analyzed control data. S.F., A.M., D.S., C.K., and K.B. analyzed the data. S.F., B.D., and T.W. acquired funding and supervised the experiments. S.F. wrote the manuscript, and A.M., D.S., C.K., H.J., T.W., and B.D. reviewed, edited, and approved the final version.

## 8 Acknowledgements

We thank Zed for his dedicated participation and his family for their support during this study.

## 10 Additional Information

The author(s) declare no competing interests

## References

1. Blom, J. D., Ter Meulen, B. C., Dool, J., & Ffytche, D. H. (2021). A century of prosopometamorphopsia studies. Cortex, 139, 298–308. 10.1016/j.cortex.2021.03.001

2. Herald, S. B., Almeida, J., & Duchaine, B. (2023). Face distortions in prosopometamorphopsia provide new insights into the organization of face perception. Neuropsychologia, 182, 108517. 10.1016/j.neuropsychologia.2023.108517

3. Lachmund, Dr. (1904). Ueber vereinzelt auftretende Halluzinationen bei Epileptikern. Monatsschrift Für Psychiatrie Und Neurologie, 15(6), 434–444. 10.1159/000219292

4. Critchley, M. (1953). The parietal lobes (pp. vii, 480). Williams and Wilkins.

5. Jonas, J., & Rossion, B. (2021). Intracerebral electrical stimulation to understand the neural basis of human face identity recognition. European Journal of Neuroscience, 54(1), 4197–4211. 10.1111/ejn.15235

6. Parvizi, J., Jacques, C., Foster, B. L., Withoft, N., Rangarajan, V., Weiner, K. S., & Grill-Spector, K. (2012). Electrical Stimulation of Human Fusiform Face-Selective Regions Distorts Face Perception. Journal of Neuroscience, 32(43), 14915–14920. 10.1523/JNEUROSCI.2609-12.2012

7. Rangarajan, V., Hermes, D., Foster, B. L., Weiner, K. S., Jacques, C., Grill-Spector, K., & Parvizi, J. (2014). Electrical Stimulation of the Left and Right Human Fusiform Gyrus Causes Different Effects in Conscious Face Perception. Journal of Neuroscience, 34(38), 12828–12836. 10.1523/JNEUROSCI.0527-14.2014

8. Puce, A., Allison, T., & McCarthy, G. (1999). Electrophysiological Studies of Human Face Perception. III: Effects of Top-down Processing on Face-specific Potentials. Cerebral Cortex, 9(5), 445–458. 10.1093/cercor/9.5.445

9. Castaldi, E., Piazza, M., & Iuculano, T. (2020). Learning disabilities: Developmental dyscalculia. Handbook of Clinical Neurology, 174, 61–75. 10.1016/B978-0-444-64148-9.00005-3

10. Mello, A., & Duchaine, B. (in press). What do we know about people with developmental prosopagnosia? K. Lander (Ed.), From Super-Recognisers to the Face Blind: Why Are Some People Better at Recognising Faces?

11. Peterson, R. L., & Pennington, B. F. (2015). Developmental dyslexia. Annual Review of Clinical Psychology, 11, 283–307. 10.1146/annurev-clinpsy-032814-112842

12. Blom, J. D., Sommer, I. E. C., Koops, S., & Sacks, O. W. (2014). Prosopometamorphopsia and facial hallucinations. The Lancet, 384(9958), 1998. 10.1016/S0140-6736(14)61690-1

13. Warrington, E. K. (1984). Recognition memory test. Western Psychological Services.

14. Mello, A., Stehr, D., Bujarski, K., & Duchaine, B. (2024). Visualising facial distortions in prosopometamorphopsia. Lancet (London, England), 403(10432), 1176. 10.1016/S0140-6736(24)00136-3

15. Rezlescu, C., Danaila, I., Miron, A., & Amariei, C. (2020). Chapter 13 - More time for science: Using Testable to create and share behavioral experiments faster, recruit better participants, and engage students in hands-on research. In B. L. Parkin (Ed.), Progress in Brain Research (Vol. 253, pp. 243–262). Elsevier. 10.1016/bs.pbr.2020.06.005

16. Kieseler, M.-L., Dickstein, A., Krafian, A., Li, C., & Duchaine, B. (2022). HEVA – A new basic visual processing test. Journal of Vision, 22(14), 4109. 10.1167/jov.22.14.4109

17. Ishihara, S. (1960). Tests for colour-blindness. Kanehara Shuppan Company Japan, *5th*.

18. Farnsworth, D. (1943). The Farnsworth-Munsell 100-Hue and Dichotomous Tests for Color Vision*. JOSA, 33(10), 568–578. 10.1364/JOSA.33.000568

19. Kinnear, P. R. (2002). New Farnsworth-Munsell 100 hue test norms of normal observers for each year of age 5-22 and for age decades 30-70. British Journal of Ophthalmology, 86(12), 1408–1411. 10.1136/bjo.86.12.1408

20. Marks, D. F. (1973). Visual imagery differences in the recall of pictures. British Journal of Psychology, 64, 17–24. 10.1111/j.2044-8295.1973.tb01322.x

21. Marks, D. F. (1995). New directions for mental imagery research. Journal of Mental Imagery, 19, 153–167.

22. Friedlander, K. J., Lenton, F. H., & Fine, P. A. (2022). A multifactorial model of visual imagery and its relationship to creativity and the vividness of Visual Imagery Questionnaire. Psychology of Aesthetics, Creativity, and the Arts, No Pagination Specified-No Pagination Specified. 10.1037/aca0000520

23. Keogh, R., & Pearson, J. (2018). The blind mind: No sensory visual imagery in aphantasia. Cortex; a Journal Devoted to the Study of the Nervous System and Behavior, 105, 53–60. 10.1016/j.cortex.2017.10.012

24. Crouzet, S. M., Kirchner, H., & Thorpe, S. J. (2010). Fast saccades toward faces: Face detection in just 100 ms. Journal of Vision, 10(4), 16. 10.1167/10.4.16

25. Tsao, D. Y., & Livingstone, M. S. (2008). Mechanisms of Face Perception. Annual Review of Neuroscience, 31(Volume 31, 2008), 411–437. 10.1146/annurev.neuro.30.051606.094238

26. Duchaine, B., Rezlescu, C., Garrido, L., Zhang, Y., Braga, M. V., & Susilo, T. (2023). The development of upright face perception depends on evolved orientation-specific mechanisms and experience. iScience, 107763. 10.1016/j.isci.2023.107763

27. Rezlescu, C., Chapman, A., Susilo, T., & Caramazza, A. (2016, December 8). Large inversion effects are not specific to faces and do not vary with object expertise [Working/ discussion paper]. Scientific Reports; PsyArXiv Preprints. 10.31234/osf.io/xzbe5

28. Duchaine, B., & Nakayama, K. (2006). The Cambridge Face Memory Test: Results for neurologically intact individuals and an investigation of its validity using inverted face stimuli and prosopagnosic participants. Neuropsychologia, 44(4), 576–585. 10.1016/j.neuropsychologia.2005.07.001

29. Duchaine, B., & Nakayama, K. (2005). Dissociations of Face and Object Recognition in Developmental Prosopagnosia. Journal of Cognitive Neuroscience, 17(2), 249–261. 10.1162/0898929053124857

30. Kieseler, M.-L., & Duchaine, B. (2023). Persistent prosopagnosia following COVID-19. Cortex, 162, 56–64. 10.1016/j.cortex.2023.01.012

31. Duchaine, B., Yovel, G., & Nakayama, K. (2007). No global processing deficit in the Navon task in 14 developmental prosopagnosics. Social Cognitive and Affective Neuroscience, 2(2), 104–113. 10.1093/scan/nsm003

32. Thompson, P. (1980). Margaret Thatcher: A new illusion. Perception, 9(4), 483–484. 10.1068/p090483

33. Dennett, H. W., McKone, E., Tavashmi, R., Hall, A., Pidcock, M., Edwards, M., & Duchaine, B. (2012). The Cambridge Car Memory Test: A task matched in format to the Cambridge Face Memory Test, with norms, reliability, sex differences, dissociations from face memory, and expertise effects. Behavior Research Methods, 44(2), 587–605. 10.3758/s13428-011-0160-2

34. Garrido, L., Duchaine, B., & Nakayama, K. (2008). Face detection in normal and prosopagnosic individuals. Journal of Neuropsychology, 2(1), 119–140. 10.1348/174866407X246843

35. Dalrymple, K. A., Davies-Thompson, J., Oruc, I., Handy, T. C., Barton, J. J. S., & Duchaine, B. (2014). Spontaneous perceptual facial distortions correlate with ventral occipitotemporal activity. Neuropsychologia, 59, 179–191. 10.1016/j.neuropsychologia.2014.05.005

36. DeBruine, L., & Jones, B. (2017). Face Research Lab London Set [Dataset]. figshare. 10.6084/m9.figshare.5047666.v5

37. Heutink, J., Brouwer, W. H., Kums, E., Young, A., & Bouma, A. (2012). When family looks strange and strangers look normal: A case of impaired face perception and recognition after stroke. Neurocase, 18(1), 39–49. 10.1080/13554794.2010.547510

38. Ebner, N. C., Riediger, M., & Lindenberger, U. (2010). FACES - A database of facial expressions in young, middle-aged, and older women and men: Development and validation. Behavior Research Methods, 42(1), 351–362. 10.3758/BRM.42.1.351

39. Egger, H. L., Pine, D. S., Nelson, E., Leibenluft, E., Ernst, M., Towbin, K. E., & Angold, A. (2011). The NIMH Child Emotional Faces Picture Set (NIMH-ChEFS): A new set of children’s facial emotion stimuli. International Journal of Methods in Psychiatric Research, 20(3), 145–156. 10.1002/mpr.343

40. Ma, D. S., Correll, J., & Wittenbrink, B. (2015). The Chicago face database: A free stimulus set of faces and norming data. Behavior Research Methods, 47(4), 1122–1135. 10.3758/s13428-014-0532-5

41. Fischl, B., & Dale, A. M. (2000). Measuring the thickness of the human cerebral cortex from magnetic resonance images. Proceedings of the National Academy of Sciences, 97(20), 11050–11055. 10.1073/pnas.200033797

42. Cox, R. W. (1996). AFNI: Software for Analysis and Visualization of Functional Magnetic Resonance Neuroimages. Computers and Biomedical Research, 29(3), 162–173. 10.1006/cbmr.1996.0014

43. Reynolds, R. C., Taylor, P. A., & Glen, D. R. (2023). Quality control practices in FMRI analysis: Philosophy, methods and examples using AFNI. Frontiers in Neuroscience, 16, 1073800. 10.3389/fnins.2022.1073800

44. Jiahui, G., Yang, H., & Duchaine, B. (2018). Developmental prosopagnosics have widespread selectivity reductions across category-selective visual cortex. Proceedings of the National Academy of Sciences, 115(28). 10.1073/pnas.1802246115

45. Norman-Haignere, S., Kanwisher, N., & McDermott, J. H. (2013). Cortical pitch regions in humans respond primarily to resolved harmonics and are located in specific tonotopic regions of anterior auditory cortex. The Journal of Neuroscience: The Official Journal of the Society for Neuroscience, 33(50), 19451–19469. 10.1523/JNEUROSCI.2880-13.2013

46. Crawford, J. R., Garthwaite, P. H., & Ryan, K. (2011). Comparing a single case to a control sample: Testing for neuropsychological deficits and dissociations in the presence of covariates. Cortex, 47(10), 1166–1178. 10.1016/j.cortex.2011.02.017

47. Duchaine, B., & Yovel, G. (2015). A Revised Neural Framework for Face Processing. Annual Review of Vision Science, 1(1), 393–416. 10.1146/annurev-vision-082114-035518

48. Rosenke, M., van Hoof, R., van den Hurk, J., Grill-Spector, K., & Goebel, R. (2021). A Probabilistic Functional Atlas of Human Occipito-Temporal Visual Cortex. Cerebral Cortex, 31(1), 603–619. 10.1093/cercor/bhaa246

49. Weiner, K. S., Golarai, G., Caspers, J., Chuapoco, M. R., Mohlberg, H., Zilles, K., Amunts, K., & Grill-Spector, K. (2014). The mid-fusiform sulcus: A landmark identifying both cytoarchitectonic and functional divisions of human ventral temporal cortex. NeuroImage, 84, 453–465. 10.1016/j.neuroimage.2013.08.068

50. Jung, H., Amini, M., Hunt, B. J., Murphy, E. I., Sadil, P., Halchenko, Y. O., Petre, B., Miao, Z., Kragel, P. A., Han, X., Heilicher, M. O., Sun, M., Collins, O. G., Lindquist, M. A., & Wager, T. D. (2025). Spacetop: A multimodal fMRI dataset unifying naturalistic processes with a rich array of experimental tasks. Scientific Data, 12(1), 1465. 10.1038/s41597-025-05154-x

51. Mori, S., Oishi, K., Jiang, H., Jiang, L., Li, X., Akhter, K., Hua, K., Faria, A. V., Mahmood, A., Woods, R., Toga, A. W., Pike, G. B., Neto, P. R., Evans, A., Zhang, J., Huang, H., Miller, M. I., van Zijl, P., & Mazziotta, J. (2008). Stereotaxic white matter atlas based on diffusion tensor imaging in an ICBM template. NeuroImage, 40(2), 570–582. 10.1016/j.neuroimage.2007.12.035

52. Oishi, K., Zilles, K., Amunts, K., Faria, A., Jiang, H., Li, X., Akhter, K., Hua, K., Woods, R., Toga, A. W., Pike, G. B., Rosa-Neto, P., Evans, A., Zhang, J., Huang, H., Miller, M. I., van Zijl, P. C. M., Mazziotta, J., & Mori, S. (2008). Human brain white matter atlas: Identification and assignment of common anatomical structures in superficial white matter. NeuroImage, 43(3), 447–457. 10.1016/j.neuroimage.2008.07.009

53. Crawford, J. R., Garthwaite, P. H., & Porter, S. (2010). Point and interval estimates of effect sizes for the case-controls design in neuropsychology: Rationale, methods, implementations, and proposed reporting standards. Cognitive Neuropsychology, 27(3), 245–260. 10.1080/02643294.2010.513967

54. Benjamini, Y., & Hochberg, Y. (1995). Controlling the False Discovery Rate: A Practical and Powerful Approach to Multiple Testing. Journal of the Royal Statistical Society: Series B (Methodological), 57(1), 289–300. 10.1111/j.2517-6161.1995.tb02031.x

55. Duerden, E. G., Card, D., Lax, I. D., Donner, E. J., & Taylor, M. J. (2013). Alterations in frontostriatal pathways in children born very preterm. Developmental Medicine & Child Neurology, 55(10), 952–958. 10.1111/dmcn.12198

56. Kanel, D., Vanes, L. D., Pecheva, D., Hadaya, L., Falconer, S., Counsell, S. J., Edwards, D. A., & Nosarti, C. (2021). Neonatal White Matter Microstructure and Emotional Development during the Preschool Years in Children Who Were Born Very Preterm. eNeuro, 8(5), ENEURO.0546-20.2021. 10.1523/ENEURO.0546-20.2021

57. Mouka, V., Drougia, A., Xydis, V. G., Astrakas, L. G., Zikou, A. K., Kosta, P., Andronikou, S., & Argyropoulou, M. I. (2019). Functional and structural connectivity of the brain in very preterm babies: Relationship with gestational age and body and brain growth. Pediatric Radiology, 49(8), 1078–1084. 10.1007/s00247-019-04412-6

58. Mullen, K. M., Vohr, B. R., Katz, K. H., Schneider, K. C., Lacadie, C., Hampson, M., Makuch, R. W., Reiss, A. L., Constable, R. T., & Ment, L. R. (2011). Preterm birth results in alterations in neural connectivity at age 16 years. NeuroImage, 54(4), 2563–2570. 10.1016/j.neuroimage.2010.11.019

59. Vollmer, B., Lundequist, A., Mårtensson, G., Nagy, Z., Lagercrantz, H., Smedler, A.-C., & Forssberg, H. (2017). Correlation between white matter microstructure and executive functions suggests early developmental influence on long fibre tracts in preterm born adolescents. PLOS ONE, 12(6), e0178893. 10.1371/journal.pone.0178893

60. Young, J. M., Vandewouw, M. M., Morgan, B. R., Smith, M. L., Sled, J. G., & Taylor, M. J. (2018). Altered white matter development in children born very preterm. Brain Structure and Function, 223(5), 2129–2141. 10.1007/s00429-018-1614-4

61. Mello, A., Garside, D., Stehr, D., Bujarski, K., Conway, B., & Duchaine, B. (2024). Color robustly affects the intensity of facial distortions in two cases of prosopometamorphopsia. Journal of Vision, 24(10), 659. 10.1167/jov.24.10.659

62. Mello, A., & Duchaine, B. (2025). What do we know about people with developmental prosopagnosia? PsyArXiv. 10.31234/osf.io/kep3t_v1

63. Sugahara, Y., Iizuka, C., Doi, K., Matsuzaki, K., & Nagaoka, M. (2022). False recognition/misidentification of unfamiliar person after cerebral infarction: A case report. Cortex, 147, 185–193. 10.1016/j.cortex.2021.12.005

64. Landis, T., Cummings, J. L., Christen, L., Bogen, J. E., & Imhof, H.-G. (1986). Are Unilateral Right Posterior Cerebral Lesions Sufficient to Cause Prosopagnosia? Clinical and Radiological Findings in Six Additional Patients. Cortex, 22(2), 243–252. 10.1016/S0010-9452(86)80048-X

65. Jonas, J., Brissart, H., Hossu, G., Colnat-Coulbois, S., Vignal, J.-P., Rossion, B., & Maillard, L. (2018). A face identity hallucination (palinopsia) generated by intracerebral stimulation of the face-selective right lateral fusiform cortex. Cortex, 99, 296–310. 10.1016/j.cortex.2017.11.022

66. Hwang, J. Y., Ha, S. W., Cho, E. K., Han, J. H., Lee, S. H., Lee, S. Y., & Kim, D. E. (2012). A Case of Prosopometamorphopsia Restricted to the Nose and Mouth with Right Medial Temporooccipital Lobe Infarction that Included the Fusiform Face Area. *Journal of Clinical Neurology (Seoul*, Korea), 8(4), 311–313. 10.3988/jcn.2012.8.4.311

67. Goodale, M. A., & Milner, A. D. (1992). Separate visual pathways for perception and action. Trends in Neurosciences, 15(1), 20–25. 10.1016/0166-2236(92)90344-8

68. Jonas, J., Descoins, M., Koessler, L., Colnat-Coulbois, S., Sauvée, M., Guye, M., Vignal, J.-P., Vespignani, H., Rossion, B., & Maillard, L. (2012). Focal electrical intracerebral stimulation of a face-sensitive area causes transient prosopagnosia. Neuroscience, 222, 281–288. 10.1016/j.neuroscience.2012.07.021

69. Jonas, J., Rossion, B., Krieg, J., Koessler, L., Colnat-Coulbois, S., Vespignani, H., Jacques, C., Vignal, J.-P., Brissart, H., & Maillard, L. (2014). Intracerebral electrical stimulation of a face-selective area in the right inferior occipital cortex impairs individual face discrimination. NeuroImage, 99, 487–497. 10.1016/j.neuroimage.2014.06.017

70. Mundel, T., Milton, J. G., Dimitrov, A., Wilson, H. W., Pelizzari, C., Uftring, S., Torres, I., Erickson, R. K., Spire, J.-P., & Towle, V. L. (2003). Transient Inability to Distinguish Between Faces: Electrophysiologic Studies. Journal of Clinical Neurophysiology, 20(2), 102. https://journals.lww.com/clinicalneurophys/abstract/2003/04000/transient_inability_to_distinguish_between_faces_.3.aspx

71. Schalk, G., Kapeller, C., Guger, C., Ogawa, H., Hiroshima, S., Lafer-Sousa, R., Saygin, Z. M., Kamada, K., & Kanwisher, N. (2017). Facephenes and rainbows: Causal evidence for functional and anatomical specificity of face and color processing in the human brain. Proceedings of the National Academy of Sciences, 114(46), 12285–12290. 10.1073/pnas.1713447114

72. Schrouff, J., Raccah, O., Baek, S., Rangarajan, V., Salehi, S., Mourão-Miranda, J., Helili, Z., Daitch, A. L., & Parvizi, J. (2020). Fast temporal dynamics and causal relevance of face processing in the human temporal cortex. Nature Communications, 11(1), 656. 10.1038/s41467-020-14432-8

73. Jonas, J., Rossion, B., Brissart, H., Frismand, S., Jacques, C., Hossu, G., Colnat-Coulbois, S., Vespignani, H., Vignal, J.-P., & Maillard, L. (2015). Beyond the core face-processing network: Intracerebral stimulation of a face-selective area in the right anterior fusiform gyrus elicits transient prosopagnosia. Cortex, 72, 140–155. 10.1016/j.cortex.2015.05.026

74. Catani, M., & Thiebautdeschotten, M. (2008). A diffusion tensor imaging tractography atlas for virtual in vivo dissections. Cortex, 44(8), 1105–1132. 10.1016/j.cortex.2008.05.004

75. Conner, A. K., Briggs, R. G., Sali, G., Rahimi, M., Baker, C. M., Burks, J. D., Glenn, C. A., Battiste, J. D., & Sughrue, M. E. (2018). A Connectomic Atlas of the Human Cerebrum—Chapter 13: Tractographic Description of the Inferior Fronto-Occipital Fasciculus. Operative Neurosurgery, 15(suppl_1), S436. 10.1093/ons/opy267

76. Motomura, K., Fujii, M., Maesawa, S., Kuramitsu, S., Natsume, A., & Wakabayashi, T. (2014). Association of dorsal inferior frontooccipital fasciculus fibers in the deep parietal lobe with both reading and writing processes: A brain mapping study: Case report. Journal of Neurosurgery, 121(1), 142–148. 10.3171/2014.2.JNS131234

77. Ortibus, E., Verhoeven, J., Sunaert, S., Casteels, I., De Cock, P., & Lagae, L. (2012). Integrity of the inferior longitudinal fasciculus and impaired object recognition in children: A diffusion tensor imaging study. Developmental Medicine & Child Neurology, 54(1), 38–43. 10.1111/j.1469-8749.2011.04147.x

78. Taddei, M., Tettamanti, M., Zanoni, A., Cappa, S., & Battaglia, M. (2012). Brain white matter organisation in adolescence is related to childhood cerebral responses to facial expressions and harm avoidance. NeuroImage, 61(4), 1394–1401. 10.1016/j.neuroimage.2012.03.062

79. Philippi, C. L., Mehta, S., Grabowski, T., Adolphs, R., & Rudrauf, D. (2009). Damage to Association Fiber Tracts Impairs Recognition of the Facial Expression of Emotion. Journal of Neuroscience, 29(48), 15089–15099. 10.1523/JNEUROSCI.0796-09.2009

