## SupplementalMaterials for "The first comprehensive case study of early-emerging prosopometamorphopsia"

### 1 Supplementary Materials

| **Table S1.** Neuropsychological, Cognitive, and Behavioral Measures of Zed and Controls | | | | | | | | |
| --- | --- | --- | --- | --- | --- | --- | --- | --- |
| **Test** | **Category** | **Zed’s score** | **Mean Control Score (SD)** | **Modified *t*-value** | ***p*-value (two-tailed)** | **Mean Control Age (SD)** | **Sample Size** | **Control Data Source** |
| Visual Vividness Imagery Questionnaire (VVIQ) | Low-level sensory visual imagery | 0.8 | 3.35 (0.93) | -2.726 | **0.008** | 27.8 (13.97) | 87 | Friedlander et al., 2024 |
| Hanover Early Vision Assessment (HEVA) | Low-level visual processing | 83% | 74.6% (10.1%) | 0.822 | 0.413 | 18.9 (1) | 117 | Kieseler et al., 2022 |
| Farnsworth Munsell 100-Hue Test for Color Vision | Color discrimination ability | 164 | 65 (14.7) | -6.621 | **<0.001** | 15 (0) | 29 | Kinnear, 2002 |
| Cambridge Face Memory Test (CFMT) | Face Recognition | 56% | 80.4% (11%) | -2.196 | **0.016** | 20.2 (1.8) | 50 | Duchaine & Nakayama, 2006 |
| Faces Old/New Test | Face Recognition | A’ 0.81 | A’ 0.965 (0.02) | -7.532 | **< 0.001** | 27.8 (N/A) | 17 | Duchaine et al., 2006 |
| Famous Faces Test | Face Recognition | 44% | 74.3% (15.1%) | -1.943 | 0.072 | 18 (0) | 15 | Prolific |
| Doppelganger Test | Face Recognition | 71% | 78.6% (10.32%) | -0.713 | 0.487 | 18 (0) | 15 | Prolific |
| Cambridge Face Perception Identity Test (CFPT-Identity) | Upright Face Perception | 54% | 74% (8.47%) | -2.267 | **0.040** | 22.4 (2.23) | 15 | Duchaine et al., 2007 |
|  | Inverted Face Perception | 39% | 54.9% (6.80%) | -2.273 | **0.039** | 22.4 (2.23) | 15 |  |
| Cambridge Face Perception Test; Age Test (CFPT-Age) | Face Perception | 72% | 76.3% (9.28%) | -0.405 | 0.688 | 30-67 (N/A) | 30 | Chatterjee & Nakayama, 2012 |
| Cambridge Face Perception Test; Sex (CFPT-Sex) | Face Perception | 78% | 76% (11%) | 0.148 | 0.884 | 30-67 (N/A) | 30 | Chatterjee & Nakayama, 2012 |
| Face Matching | Upright Face Identity Perception | 78% | 78.7% (11.6%) | -0.013 | 0.919 | 36.1 (N/A) | 63 | Rezlescu et al., 2024 |
|  | Inverted Face Identity Perception | 45% | 53.6% (13.5%) | -0.632 | 0.530 | 36.1 (N/A) | 63 |  |
| Thatcher Test | Upright Face Perception | 97% | 95.8% (11.5%) | 0.074 | 0.942 | 39.41 (3.7) | 22 | Duchaine et al., 2023 |
|  | Inverted Face Perception | 50% | 51% (10%) | -0.098 | 0.923 | 39.41 (3.7) | 22 |  |
| Face Mooney Forced-Choice Test | Upright Face Detection | 82% | 87.5% (5.65%) | -0.946 | 0.355 | 39.41 (3.7) | 22 |  |
|  | Inverted Face Detection | 64% | 61.22% (10.91%) | 0.259 | 0.798 | 39.41 (3.7) | 22 |  |
| Two-Tone Forced-Choice Test | Upright Face Detection | 82% | 95.8% (5.54%) | -2.436 | **0.024** | 39.41 (3.7) | 22 |  |
| Voices Old/New Test | Voice Recognition | A’ 0.78 | A’ 0.76 (0.14) | 0.136 | 0.895 | 28 (N/A) | 10 | Kieseler & Duchaine, 2023 |
| Voices Learning 1 | Voice Recognition | 78% | 75% (10.9%) | 0.262 | 0.799 | 28 (N/A) | 10 |  |
| Voices Learning 2 | Voice Recognition | 65% | 70.5% (11.1%) | -0.472 | 0.648 | 28 (N/A) | 10 |  |
| Voices Learning 3 | Voice Recognition | 82% | 75.2% (9.5%) | 0.652 | 0.530 | 28 (N/A) | 10 |  |
| Voice Recognition | Voice Recognition | 25% | 28.8% (16.6%) | -0.218 | 0.832 | 28 (N/A) | 10 |  |
| Cars Old/New Test | Car Recognition | A’ 0.85 | A’ 0.94 (0.04) | -2.24 | **0.040** | 27.8 (N/A) | 17 | Duchaine et al., 2006 |
| Cambridge Car Memory Test (CCMT) | Car Recognition | 56% | 70.6% (9.93%) | -1.506 | 0.134 | 20.63 (N/A) | 153 | Dennett et al., 2012 |
| Car Matching Test | Upright Car Perception | 78% | 76.5% (10.1%) | 0.098 | 0.922 | 36.1 (10.1) | 63 | Rezlescu et al., 2024 |
|  | Inverted Car Perception | 55% | 53.2% (11.1%) | 0.161 | 0.873 | 36.1 (10.1) | 63 |  |
| Car Mooney Forced-Choice Test | Upright Car Detection | 90% | 75.1% (11.7%) | 1.242 | 0.219 | 36.1 (10.1) | 63 | Rezlescu et al., 2024 |
|  | Inverted Car Detection | 64% | 46.7% (9.7%) | 1.78 | 0.080 | 36.1 (10.1) | 63 |  |

**Supplementary Table S1. Neuropsychological, Cognitive, and Behavioral Measures of Zed and Controls.** All statistical analyses are modified via Crawford’s t-test for single cases (Crawford et al., 2011).

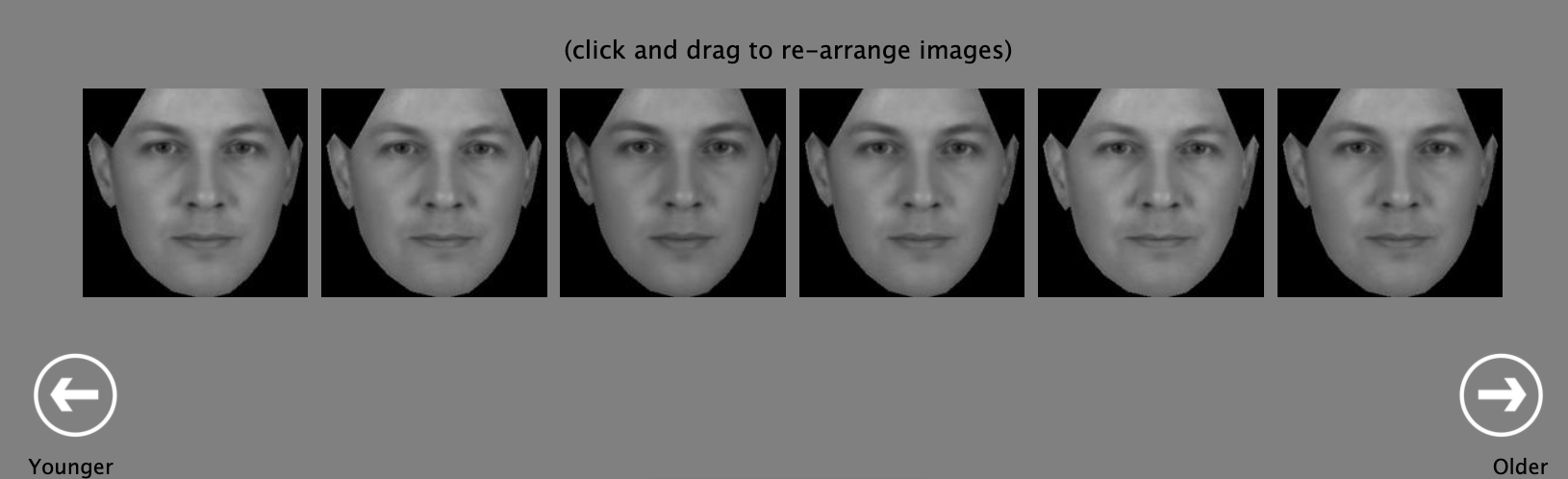

**Supplementary Figure S1. Cambridge Face Perception Test Age Stimuli.** Participants are asked to drag the faces to sort them by their perceived age from “Younger” to “Older”. There is a 40 second timer for each set of faces.

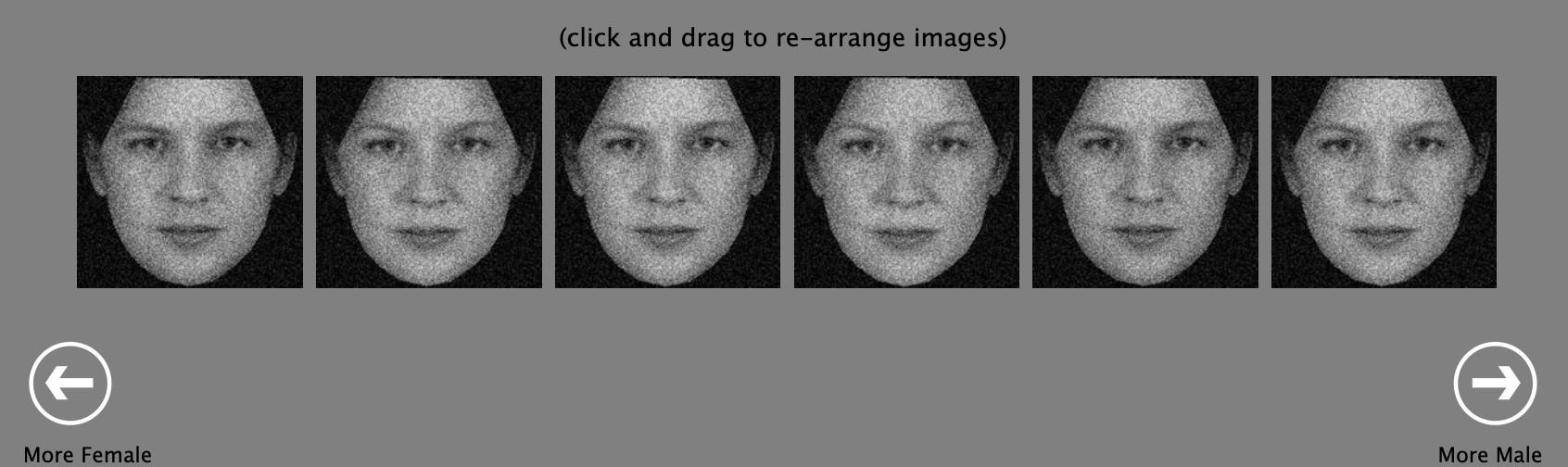

**Supplementary Figure S2. Cambridge Face Perception Test Sex Stimuli.** Participants are asked to drag the faces to sort them by their perceived sex from “More Female” to “More Male”. There is a 40 second timer for each set of faces.

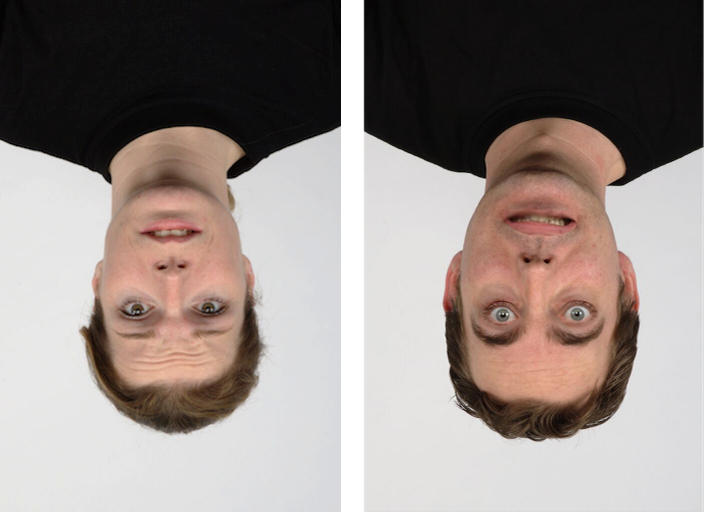

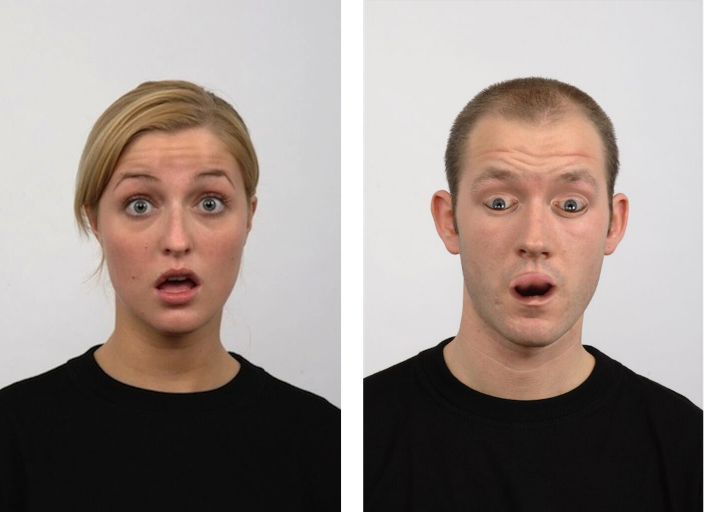

**Supplementary Figure S3.** **Thatcher Test Stimuli.** Thatcherized stimuli are created by rotating the eyes and mouth 180 degrees while the rest of the face is unchanged. **(Left)** The upright images are shown in a pair on the left. The normal face is shown on the left side of the pair, and the Thatcherized face is shown on the right side of the pair. **(Right)** The inverted images are shown in a pair on the right. The normal face is shown on the left side of the pair, and the Thatcherized face is shown on the right side of the pair.

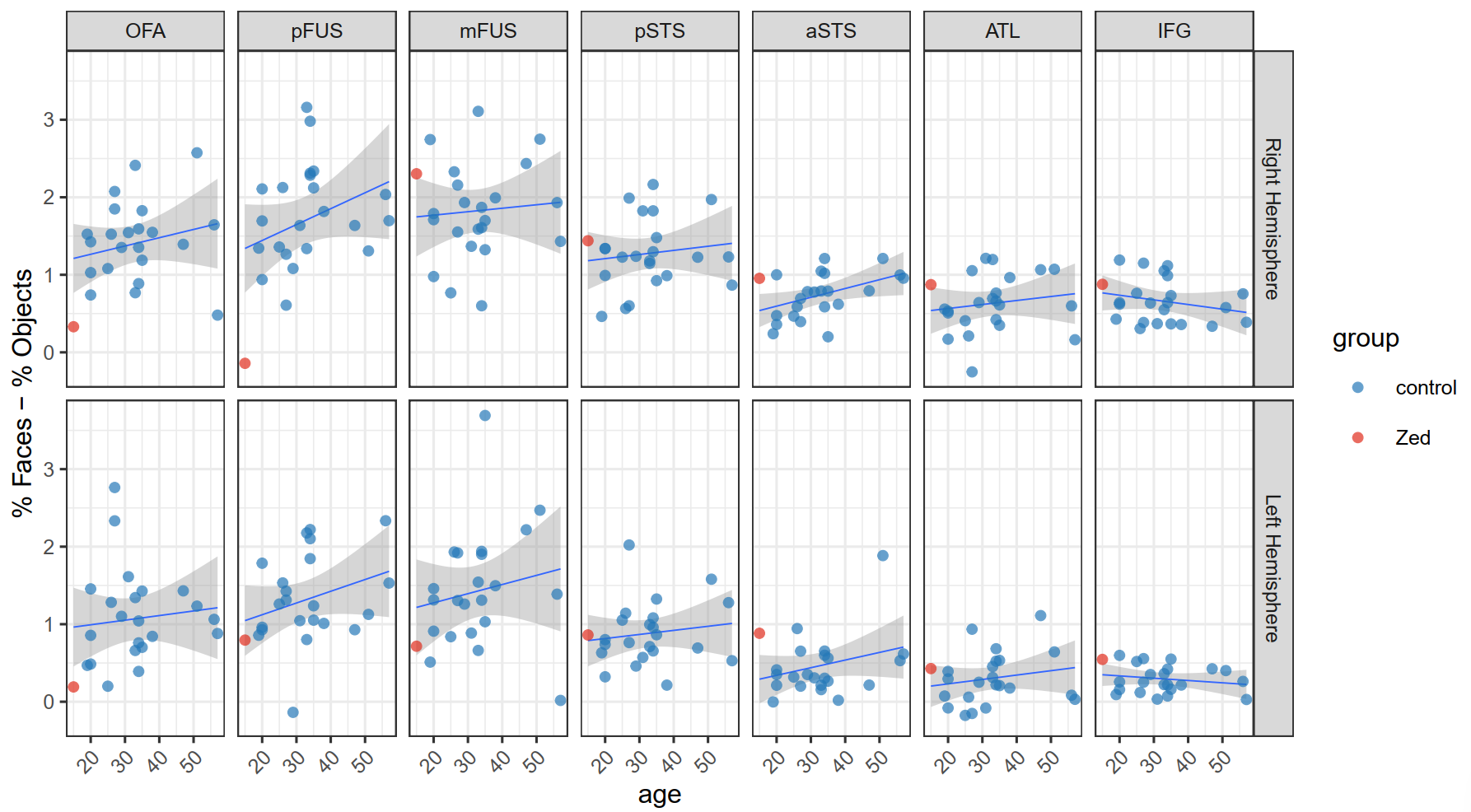

**Supplementary Figure S4. Face selectivity (faces-objects) related to age in Zed vs. Controls.** Face selectivity expressed as percent signal change to faces minus percent signal change to objects for Zed and each control by ROI and hemisphere, with regression slopes relating age to face selectivity in the controls. Shading around the lines indicates the standard error of the mean.

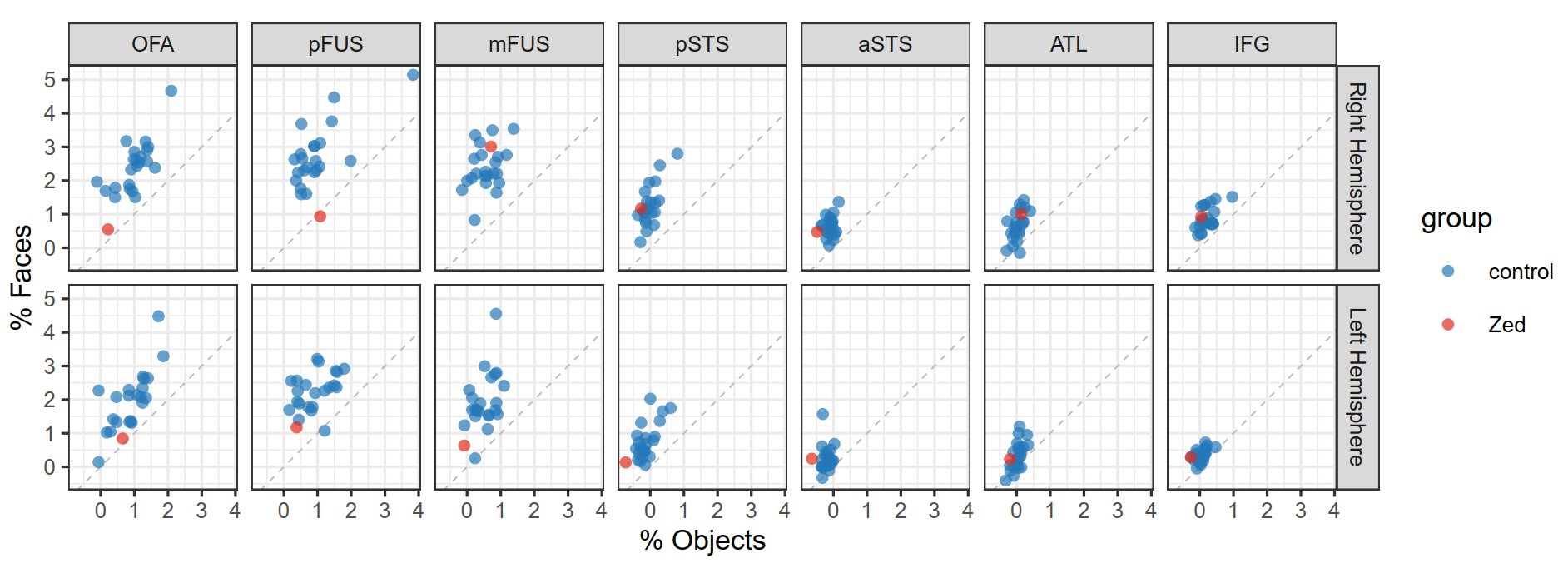

**Supplementary Figure S5. Face selectivity (faces-objects) in Zed vs. Controls.** The face-selectivity values displayed elsewhere were generated by subtracting the response to objects from the response to faces. Because atypical face-selectivity values can result from an atypical response to faces, objects, or both categories, here we show percent signal change to objects plotted against percent signal change to faces. Points are individual participants, and the dashed line indicates the line of equivalence.

| Table S2. Zed’s Multiview Face Distortion Assessment (MFDA) Results | | |
| --- | --- | --- |
| Presentation | **Stimulus Type** | **Mean Distortion Severity Rating** |
| Visible Facial Features | Nose | 0.833 |
|  | Left Eye | 1 |
|  | Right Eye | 1.5 |
|  | Mouth | 1.167 |
|  | Eyes | 1.916 |
|  | Bottom Half | 2.25 |
|  | Top Half | 2.083 |
|  | Right Half | 2.083 |
|  | Left Half | 2.25 |
|  | Full | 2.333 |
| Picture-plane orientation | 0 degrees | 2.75 |
|  | 45 degrees | 2.92 |
|  | 90 degrees | 3.50 |
|  | 135 degrees | 3.17 |
|  | 180 degrees | 2.916 |
|  | 225 degrees | 2.83 |
|  | 270 degrees | 3 |
|  | 315 degrees | 3.08 |
| Visual field position | Upper Right | 3.833 |
|  | Lower Right | 4.25 |
|  | Center | 2.916 |
|  | Upper Left | 4.08 |
|  | Lower Left | 4.42 |
| Face viewpoint | 175px by 175px | 2.42 |
|  | 350px by 350px | 2.25 |
|  | 525px by 525px | 1.92 |
|  | 700px by 700px | 2 |
| Visual angle | Frontal | 2.417 |
|  | Left 35 degree | 1.92 |
|  | Left Profile | 2.5 |
|  | Right 35 degree | 2.5 |
|  | Right Profile | 2.58 |

**Supplementary Table S2. Zed’s Multiview Face Distortion Assessment (MFDA) Results.** Each dimension includes between 48 and 120 photographs. After viewing each face, participants rate the severity of their distortions on a Likert-type scale from 0 to 6 (0 = No distortion, 6 = Extreme distortion).

| **Table S3.** Zed’s MFDA Position Condition Results | | |
| --- | --- | --- |
| **Stimulus Type** | **Mean Distortion Severity Rating** | **SEM Distortion Severity Rating** |
| Upper Right | 3.83 | 0.11 |
| Lower Right | 4.25 | 0.22 |
| Center | 2.92 | 0.15 |
| Upper Left | 4.08 | 0.19 |
| Lower Left | 4.42 | 0.19 |

**Supplementary Table S3.** **Zed’s Multiview Face Distortion Assessment Position Condition Mean and SEM Distortion Severity Ratings.** Zed experienced a 1.23-point increase in average distortion intensity rating when presented with faces in his periphery.

| **Table S4.** Zed’s Distortion Frequency in MFDA Visible Facial Features Condition | | | | |
| --- | --- | --- | --- | --- |
| **Stimulus Type** | **Mean Distortion Severity Rating** | **SEM Distortion Severity Rating** | **# of Stimuli without Distortion** | **% of Stimuli without Distortion** |
| Nose | 0.833 | 0.241 | 5/6 | 83.3% |
| Left Eye | 1 | 0.246 | 4/6 | 66.7% |
| Right Eye | 1.5 | 0.230 | 1/6 | 16.7% |
| Mouth | 1.167 | 0.207 | 2/6 | 33.3% |
| Eyes | 1.916 | 0.193 | 0/6 | 0% |
| Bottom Half | 2.25 | 0.179 | 0/6 | 0% |
| Top Half | 2.083 | 0.149 | 0/6 | 0% |
| Right Half | 2.083 | 0.193 | 0/6 | 0% |
| Left Half | 2.25 | 0.179 | 0/6 | 0% |
| Full | 2.333 | 0.188 | 0/6 | 0% |

**Supplementary Table S4.** **Zed’s Distortion Frequency in the Multiview Face Distortion Assessment Visible Facial Features Condition.** Zed’s mean distortion severity rating decreases when perceiving individual features. This effect is primarily driven by ratings of 0 when perceiving individual features.

### 2 References

Chatterjee, G., & Nakayama, K. (2012). Normal facial age and gender perception in developmental prosopagnosia. *Cognitive Neuropsychology*, *29*(5–6), 482–502. https://doi.org/10.1080/02643294.2012.756809

Crawford, J. R., Garthwaite, P. H., & Ryan, K. (2011). Comparing a single case to a control sample: Testing for neuropsychological deficits and dissociations in the presence of covariates. *Cortex*, *47*(10), 1166–1178. https://doi.org/10.1016/j.cortex.2011.02.017

Dennett, H. W., McKone, E., Tavashmi, R., Hall, A., Pidcock, M., Edwards, M., & Duchaine, B. (2012). The Cambridge Car Memory Test: A task matched in format to the Cambridge Face Memory Test, with norms, reliability, sex differences, dissociations from face memory, and expertise effects. *Behavior Research Methods*, *44*(2), 587–605. https://doi.org/10.3758/s13428-011-0160-2

Duchaine, B., & Nakayama, K. (2006). The Cambridge Face Memory Test: Results for neurologically intact individuals and an investigation of its validity using inverted face stimuli and prosopagnosic participants. *Neuropsychologia*, *44*(4), 576–585. https://doi.org/10.1016/j.neuropsychologia.2005.07.001

Duchaine, B., Rezlescu, C., Garrido, L., Zhang, Y., Braga, M. V., & Susilo, T. (2023). The development of upright face perception depends on evolved orientation-specific mechanisms and experience. *iScience*, 107763. https://doi.org/10.1016/j.isci.2023.107763

Duchaine, B., Yovel, G., Butterworth, E. J., & Nakayama, K. (2006). Prosopagnosia as an impairment to face-specific mechanisms: Elimination of the alternative hypotheses in a developmental case. *Cognitive Neuropsychology*, *23*(5), 714–747. https://doi.org/10.1080/02643290500441296

Duchaine, B., Yovel, G., & Nakayama, K. (2007). No global processing deficit in the Navon task in 14 developmental prosopagnosics. *Social Cognitive and Affective Neuroscience*, *2*(2), 104–113. https://doi.org/10.1093/scan/nsm003

Friedlander, K. J., Lenton, F. H., & Fine, P. A. (2022). A multifactorial model of visual imagery and its relationship to creativity and the vividness of Visual Imagery Questionnaire. *Psychology of Aesthetics, Creativity, and the Arts*, No Pagination Specified-No Pagination Specified. https://doi.org/10.1037/aca0000520

Kieseler, M.-L., Dickstein, A., Krafian, A., Li, C., & Duchaine, B. (2022). HEVA – A new basic visual processing test. *Journal of Vision*, *22*(14), 4109. https://doi.org/10.1167/jov.22.14.4109

Kieseler, M.-L., & Duchaine, B. (2023). Persistent prosopagnosia following COVID-19. *Cortex*, *162*, 56–64. https://doi.org/10.1016/j.cortex.2023.01.012

Kinnear, P. R. (2002). New Farnsworth-Munsell 100 hue test norms of normal observers for each year of age 5-22 and for age decades 30-70. *British Journal of Ophthalmology*, *86*(12), 1408–1411. https://doi.org/10.1136/bjo.86.12.1408

Rezlescu, C., Chapman, A., Susilo, T., & Caramazza, A. (2024). *Large inversion effects are not specific to faces and do not vary with object expertise*. https://doi.org/10.31234/osf.io/xzbe5
